# A community-driven resource for genomic epidemiology and antimicrobial resistance prediction of *Neisseria gonorrhoeae* at Pathogenwatch

**DOI:** 10.1101/2020.07.03.186726

**Authors:** Leonor Sánchez-Busó, Corin A. Yeats, Benjamin Taylor, Richard J. Goater, Anthony Underwood, Khalil Abudahab, Silvia Argimón, Kevin C. Ma, Tatum D. Mortimer, Daniel Golparian, Michelle J. Cole, Yonatan H. Grad, Irene Martin, Brian H. Raphael, William M. Shafer, Gianfranco Spiteri, Katy Town, Teodora Wi, Simon R. Harris, Magnus Unemo, David M. Aanensen

## Abstract

**Background:** Antimicrobial resistant (AMR) *Neisseria gonorrhoeae* is an urgent threat to public health, as strains resistant to at least one of the two last line antibiotics used in empiric therapy of gonorrhoea, ceftriaxone and azithromycin, have spread internationally. Whole genome sequencing (WGS) data can be used to identify new AMR clones, transmission networks and inform the development of point-of-care tests for antimicrobial susceptibility, novel antimicrobials and vaccines. Community driven tools that provide an easy access to and analysis of genomic and epidemiological data is the way forward for public health surveillance.

**Methods:** Here we present a public health focussed scheme for genomic epidemiology of *N. gonorrhoeae* at Pathogenwatch (https://pathogen.watch/ngonorrhoeae). An international advisory group of experts in epidemiology, public health, genetics and genomics of *N. gonorrhoeae* was convened to inform on the utility of current and future analytics in the platform. We implement backwards compatibility with MLST, NG-MAST and NG-STAR typing schemes as well as an exhaustive library of genetic AMR determinants linked to a genotypic prediction of resistance to eight antibiotics. A collection of over 12,000 *N. gonorrhoeae* genome sequences from public archives has been quality-checked, assembled and made public together with available metadata for contextualization.

**Results:** AMR prediction from genome data revealed specificity values over 99% for azithromycin, ciprofloxacin and ceftriaxone and sensitivity values around 99% for benzylpenicillin and tetracycline. A case study using the Pathogenwatch collection of *N. gonorrhoeae* public genomes showed the global expansion of an azithromycin resistant lineage carrying a mosaic *mtr* over at least the last 10 years, emphasizing the power of Pathogenwatch to explore and evaluate genomic epidemiology questions of public health concern.

**Conclusions:** The *N. gonorrhoeae* scheme in Pathogenwatch provides customized bioinformatic pipelines guided by expert opinion that can be adapted to public health agencies and departments with little expertise in bioinformatics and lower resourced settings with internet connection but limited computational infrastructure. The advisory group will assess and identify ongoing public health needs in the field of gonorrhoea, particularly regarding gonococcal AMR, in order to further enhance utility with modified or new analytic methods.

## Background

Antimicrobial resistance (AMR) is an urgent threat to public health. *Neisseria gonorrhoeae*, the strictly human pathogen causing the sexually-transmitted infection (STI) gonorrhoea, has developed or acquired resistance to the last-line antibiotics used in empiric therapy to treat the infection, and thus has become one of the major global priorities in order to tackle AMR. In 2017, due to the increase in AMR infections and the absence of an effective vaccine, the World Health Organization (WHO) included *N. gonorrhoeae* as a high priority pathogen in need of research and development of new antimicrobials and ideally a vaccine (1). In 2019, the Centers for Disease Control and Prevention (CDC) again included the gonococcus on the list of urgent threats in the United States (2). The most recent WHO estimates from 2016 indicate an annual global incidence of 87 million cases of gonorrhoea among adults (3, 4). Untreated cases can develop complications including an increased acquisition and transmission of HIV. In women, long-term infections can cause infertility, pelvic inflammatory disease, ectopic pregnancy, miscarriage or premature labour (5). Infections during pregnancy can transmit to newborns at birth causing eye damage that can have permanent effects on vision (6).

Strains of *N. gonorrhoeae* resistant to every recommended treatment have rapidly emerged, including resistance to penicillins, tetracyclines, fluoroquinolones, macrolides and the extended-spectrum cephalosporins (ESCs) (5-8). The current recommended treatment in many countries is a dual therapy with injectable ceftriaxone plus oral azithromycin, although reports of decreased susceptibility to ceftriaxone as well as azithromycin resistance have increased globally (7, 8). One case of failure of dual treatment was reported in 2016 in the United Kingdom (UK) (9). Additionally, in 2018 a gonococcal strain with resistance to ceftriaxone combined with high-level resistance to azithromycin was detected in both the UK and Australia (10). The transmission of a ceftriaxone-resistant clone (FC428) has been documented internationally since 2015, raising concerns about the long-term effectiveness of the current treatment in the absence of an available alternative (11). In some countries such as in Japan, China and since 2019 in the UK, a single dose of ceftriaxone 1 gram is the recommended treatment due to the increasing incidence of azithromycin resistance in *N. gonorrhoeae* and other STI pathogens such as *Mycoplasma genitalium* (12). Extensive investigations have been ongoing for years to unveil the genetic mechanisms that explain most of the observed susceptibility patterns for the main classes of antimicrobials for *N. gonorrhoeae*. For ciprofloxacin, nearly all resistant strains have the GyrA S91F amino acid alteration (13-15), however, resistance prediction from genomic data is not as straightforward for other antibiotics. Known resistance mechanisms often involve additive or suppressive effects as well as epistatic interactions that all together explain just part of the observed phenotypic resistance. For example, there is good evidence that many mosaic structures of the *penA* gene are associated with decreased susceptibility to ESCs (16, 17), however, mosaics do not explain all cases of ESC resistance, especially for ceftriaxone, and some mosaic *penA* alleles do not cause decreased susceptibility or resistance to this antibiotic (16-19). On top of these, variants that overexpress the MtrCDE efflux pump, mutations in *porB1b* that reduce drug influx and non-mosaic mutations in penicillin-binding proteins also contribute to decreased susceptibility to ESCs (20). Furthermore, mutations in the *rpoB* and *rpoD* genes, encoding subunits of the RNA polymerase, have been recently related to resistance to ESCs in clinical *N. gonorrhoeae* isolates (21). Mutations in the 23S rRNA gene (A2045G and C2597T in *N. gonorrhoeae* nomenclature, coordinates from the WHO 2016 reference panel (22), A2059G and C2611T in *Escherichia coli*) are frequently associated with azithromycin resistance, as do variants in *mtrR* or its promoter that increase the expression of the MtrCDE efflux pump (5). Recently, epistatic interactions between a mosaic *mtr* promoter region and a mosaic *mtrD* gene have also been reported to increase the expression of this pump, contributing to macrolide resistance (23, 24). Mutations in *rplD* have also been associated with reduced susceptibility to this antibiotic (25) and contrarily, loss-of-function mutations in *mtrC* have been linked to increased susceptibility to several antibiotics including azithromycin (26). Thus, we can relatively confidently predict decreased susceptibility or resistance to an antimicrobial using the current known genetic mechanisms, however, phenotypic testing is still necessary to detect resistant cases caused by unknown or novel mechanisms. These inconsistencies with the genomic data will allow the discovery of these new mechanisms, which will keep improving the resistance predictions from WGS.

A myriad of methods have been used to discriminate among strains of *N. gonorrhoeae*, from phenotypic to DNA-based techniques (27), but whole genome sequencing (WGS) can provide the complete genome information of a bacterial strain. The cost of amplifying all loci of the different typing schemes via nucleic acid amplification and traditional Sanger sequencing can be more expensive than the cost of WGS of one bacterial genome in many settings. With WGS, multiple genetic AMR mechanisms as well as virulence and typing regions can be targeted simultaneously with the appropriate bioinformatic tools and pipelines. It also provides a significant improvement in resolution and accuracy over traditional molecular epidemiology and typing methods, allowing a genome-wide comparison of strains that can: identify AMR clones, outbreaks, transmission networks, national and international spread, known and novel resistance mechanisms as well as also inform on the development of point-of-care tests for antimicrobial susceptibility, novel antimicrobials and vaccines (28, 29). However, implementation of WGS for genomic surveillance poses practical challenges, especially for Low-and Middle-Income Countries (LMICs), due to the need of a major investment to acquire and maintain the required infrastructure.

WGS produces a very high volume of data that needs to be pre-processed and analysed using bioinformatics. Bioinformatics expertise is not always readily available in laboratory and public health settings, and currently there are no international standards and proficiency trials for which algorithms to use to process WGS data. There are several open-source tools specialised in each step of the pipeline as well as proprietary software containing workflows that simplify the analyses. However, these are less customizable and may not be affordable for all (30, 31). Choosing the best algorithms and parameters when analysing genomic data is not straightforward as it requires a fair knowledge of the pathogen under study and its genome diversity. Multiple databases containing genetic determinants of AMR for bacterial pathogens are available (30, 31), however, choosing which one is most complete for a particular organism frequently requires an extensive literature search. Public access web-based species-specific tools and AMR databases revised and curated by experts would be the most approachable option for both well-resourced and LMICs with a reliable internet connection. Very importantly though, the full benefits of using WGS for both molecular epidemiology and AMR prediction can only be achieved if the WGS data are linked to phenotypic data for the gonococcal isolates and, as much as feasible, clinical and epidemiological data for the patients.

Here, we present a public health focussed system to facilitate genomic epidemiology of *N. gonorrhoeae* within Pathogenwatch (https://pathogen.watch/ngonorrhoeae), which includes the latest analytics for typing, detection of genetic AMR determinants and prediction of AMR from *N. gonorrhoeae* genome data, linked to metadata where available, as well as a collection of over 12,000 gonococcal genomes from public archives for contextualization. We formed an advisory group including experts in the field of *N. gonorrhoeae* epidemiology, public health, AMR, genetics and genomics to consult on the development and design of the tool, such as the analytics and genetic AMR mechanisms to include, in order to adapt the platform for ongoing public health needs. We present this scheme as a community-steered model for genomic surveillance that can be applied to other pathogens.

## Methods

### The Pathogenwatch platform: technical summary

Pathogenwatch is a web-based platform with several different components. The main interface is a React (32) single-page application with a style based on Material Design Lite (33). Phylogenetic trees are plotted using Phylocanvas (34), maps using Leaflet (35) and networks with Sigma (36). The back end is written in Node.js and contains an API service for the user interface and four “Runner” services for the following analyses: species prediction, single-genome analyses, tree building and core genome multi-locus sequence typing (cgMLST) clustering. Docker containers are used for queuing tasks, streaming input or result files through standard input and storing JSON data from standard output. A MongoDB cluster is used for data storage and task queuing/synchronisation. Pathogenwatch shares some visualization components with Microreact (37), such as those associated with the phylogenetic tree and the map. However, Pathogenwatch includes an analytical framework which is unique to this platform.

### Generation of the N. gonorrhoeae core genome library

Pathogenwatch implements a library of core genome sequences for several supported organisms. In the case of *N. gonorrhoeae*, a core gene set was built from the 14 finished reference genomes that constitute the 2016 WHO reference strain panel (22) using the pangenome analysis tool Roary (38) as described in Harris *et al* (2018) (15). Briefly, the minimum percentage of identity for blastp was set to 97% and the resulting core genes were aligned individually using MAFFT. The resulting genes with a percentage of identity above 99% were post-processed as described in (39). Representatives for each family were selected by choosing the sequence with the fewest differences to the others on average and searched using tblastn (percentage of identity >= 80%, E-value <= 1e-35) against the 14 high quality reference genomes. Families without a complete match in every reference (100% coverage) or had multiple matches were removed. Overlapping genes from each reference were merged into pseudocontigs and grouped by gene composition. For each family, a representative was selected as before and searched/filtered using the references as before. The final core gene set contains 1,542 sequences that span a total of 1,470,119 nucleotides (approximately 67% of a typical *N. gonorrhoeae* genome length, 2.2Mb). A BLAST database was constructed from these core segments and used to profile new assemblies.

### Profiling new assemblies

New genome assemblies can be uploaded by a user (drag and drop) or calculated from high-throughput short read data directly within Pathogenwatch using SPAdes (40) as described in (41). A taxonomy assignment step for species identification is performed on the uploaded assemblies by using Speciator (42). New assemblies are then queried against a species-specific BLAST database using blastn. For *N. gonorrhoeae*, every core loci needs to match at least 80% of its length to be considered as present. Further filtering steps are applied to remove loci that can be problematic for tree building, such as paralogs or loci with unusually large number of variant sites compared to an estimated substitution rate on the rest of the genome, as described in (43). The overall substitution rate is calculated as the number of total differences in the core library divided by the total number of nucleotides. Indels are ignored to minimise the noise that could be caused by assembly or sequencing errors. The expected number of substitutions per locus is determined by multiplying this substitution rate by the length of the representative sequence.

The number of substitutions observed for each locus between the new assembly and the reference sequence are scaled to the total number of nucleotides that match the core library, creating a pairwise score that is saved on a distance matrix and is used for Neighbour-Joining tree construction, as described in (44).

### Algorithms for sequence typing and cgMLST clustering

Alleles and sequence types (STs) for Multi-Locus Sequence Typing (MLST) (45) and cgMLST (core genome MLST, *N. gonorrhoeae* cgMLST v1.0) (46) were obtained from PubMLST (47, 48), for *N. gonorrhoeae* Multi-Antigen Sequence Typing (NG-MAST) (49) from (50) and for *N. gonorrhoeae* Sequence Typing for Antimicrobial Resistance (NG-STAR) (51) from (52) (Table 1). A search tool implemented as part of Pathogenwatch is used to make the assignments for MLST, cgMLST and NG-STAR, while NGMASTER (53) is used for NG-MAST. Briefly, exact matches to known alleles are searched for, while novel sequences are assigned a unique identifier. The combination of alleles is used to assign a ST as described in (54). Databases are regularly updated and novel alleles and STs should be submitted by the user to the corresponding schemes for designation.

**Table 1.** *N. gonorrhoeae* sequence typing schemes implemented in Pathogenwatch.

| Typing scheme* | Loci (number) | Note | Pathogenwatch implementation | References |
| --- | --- | --- | --- | --- |
| cgMLST | (N=1,649) | *N. gonorrhoeae* cgMLST v1.0 | Typing algorithm, database from PubMLST | (46-48, 89) |
| MLST | *abcZ*, *adk*, *aroE*, *fumC*, *gdh*, *pdhC*, *pgm* (N=7) | Housekeeping genes in *Neisseria* spp. | In-house typing tool, database from PubMLST | (45, 47, 48, 89) |
| NG-MAST | *porB*, *tbpB* (N=2) | Genes encoding highly-variable membrane proteins | NG-MASTER, database from NG-MAST website | (49, 50, 53) |
| NG-STAR | *penA*, *mtrR*, *porB*, *ponA*, *gyrA*, *parC*, 23S *rDNA* (N=7) | Genes involved in antimicrobial resistance | In-house typing tool, database from NG-STAR website | (51, 52, 89) |

\* Typing scheme: cgMLST = core genome Multi-Locus Sequence Typing, MLST = Multi-Locus Sequence Typing, NG-MAST = *N. gonorrhoeae* Multi-Antigen Sequence Typing, NG-STAR = *N. gonorrhoeae* Sequence Typing for Antimicrobial Resistance.

cgMLST typing information is used for clustering individual genomes with others in the Pathogenwatch database using single linkage clustering as described in (55). Users can select the clustering threshold (i.e. number of loci with differing alleles) and a network graph based on the SLINK (56) algorithm is calculated within individual genome reports.

### AMR library and detection of genetic AMR determinants

Genes and point mutations (single nucleotide polymorphisms (SNPs) and indels) were detected using Pathogenwatch AMR v2.4.9 (57). Pathogenwatch AMR also provides a prediction of AMR phenotype inferred from the combination of identified mechanisms. Genetic determinants described in the literature as involved in AMR in *N. gonorrhoeae* were collated into a library in TOML format (version 0.0.11). A test dataset containing 3,987 isolates from 13 studies (15, 18, 22, 58-67) (Additional file 1: Table S1) providing minimum inhibitory concentration (MIC) information for six antibiotics (benzylpenicillin, tetracycline, ciprofloxacin, cefixime, ceftriaxone and azithromycin) was used to benchmark and to curate this library. A validation benchmark was posteriorly run with a dataset of 1,607 isolates from 3 other publications (68-70) with MIC information for the same six antibiotics plus spectinomycin (Additional file 1: Table S1). EUCAST clinical breakpoints v9.0 (71) were used to define susceptibility (S), susceptibility with an increased exposure (I) or resistance (R) (SIR) categorical interpretations of MICs for all antibiotics except for azithromycin, for which the EUCAST epidemiological cut-off (ECOFF) was used to define non-susceptibility/resistance (ECOFF>1mg/L). As a result of the benchmark analyses, sensitivity, specificity and positive/negative predictive values (PPV/NPV) were obtained for the AMR mechanisms implemented in the library and, globally, for each of the antibiotics. Confidence intervals (95%) for these statistics were calculated using the *epi*.*tests* function in the *epiR* R package v1.0-14 (72). Individual or combined AMR mechanisms with a PPV below 15% were discarded from the library to optimise the overall predictive values. Visual representations of the observed ranges of MIC values for a particular antibiotic for each of the observed combinations of genetic AMR mechanisms on the test dataset were used to identify and assess combinations of mechanisms that have an additive or suppressive effect on AMR. These were included in the library.

As part of the accuracy testing of the AMR library, we ran the 2016 WHO *N. gonorrhoeae* reference genomes 2016 panel (n=14) through Pathogenwatch and compared the detected list of genetic AMR mechanisms with the list published in the original study (22). For the WHO U strain, a discrepancy on a mutation in *parC* was further investigated by mapping the original raw Illumina data (European Nucleotide Archive (ENA) run accession ERR449479) to the reference genome assembly (ENA genome accession LT592159.1) and visualized using Artemis (73).

In short-read assemblies, the four copies of the 23S rRNA gene are collapsed into one, thus the detection of the A2045G and C2597T mutations is dependent on the consensus bases resulting from the number of mutated copies (63, 66, 74).

### Quality check and assembly of public sequencing data

Public *N. gonorrhoeae* genomes with geolocation data were obtained from the ENA in November 2019. This list was complemented by an exhaustive literature search of studies on *N. gonorrhoeae* genomics without metadata submitted to the ENA but instead made available as supplementary information in the corresponding publications. Raw paired-end short read data from a list of 12,192 isolates was processed with the GHRU assembly pipeline v1.5.4 (75). This pipeline runs a Nextflow workflow to quality-check (QC) paired-end short read fastq files before and after filtering and trimming, assembles the data and quality-checks the resulting assembly. Results from the pipeline are provided in Additional file 2. In this pipeline, QC of short reads was performed using FastQC v0.11.8 (76). Trimming was done with Trimmomatic v0.38 (77) by cutting bases from the start and end of reads if they were below a Phred score of 25, trimming using a sliding window of size 4 and cutting once the average quality within the window fell below a Phred score of 20. Only reads with length above a third of the original minimum read length were kept for further analyses. After trimming, reads were corrected using the kmer-based approach implemented in Lighter v1.1.1 (78) with a kmer length of 32 bp and a maximum number of corrections allowed within a 20 bp window of 1. ConFindr v0.7.2 was used to assess intra- and inter-species contamination (79). Mash v2.1 (80) was applied to estimate genome size using a kmer size of 32 bp and Seqtk v1.3 (81) to down sample fastq files if the depth of coverage was above 100x. Flash v1.2.11 (82) was used to merge reads with a minimum overlap length of 20 bp and a maximum overlap of 100 bp to facilitate the subsequent assembly process. SPAdes v3.12 (40) was used for genome assembly with the --careful option selected to reduce the number of mismatches and short indels with a range of kmer lengths depending on the minimum read length. The final assemblies were quality-checked using Quast v5.0.2 (83) and ran through the species identification tool Bactinspector (84). QC conditions were assessed and summarised using Qualifyr (85).

Fastq files with poor quality in which the trimming and filtering step discarded all reads from either one or both pairs were excluded from the analyses because the assembly pipeline is optimised for paired-end data. Assemblies with an N50 below 25,000 bp, a number of contigs above 300, a total assembly length above 2.5 Mb or a percentage of contamination above 5% were also excluded.

### Metadata for public genomes

Geolocation data (mainly country), collection dates (day, month and year when available), ENA project accession and associated Pubmed ID were obtained from the ENA API for all the genomes in the pipeline (86). A manual extensive literature search was performed to identify the publications containing the selected genomes. In order to complete published studies as much as possible, extra genomes were downloaded and added to the dataset. Metadata for the final set was completed with the information contained in supplementary tables on the corresponding publications, including phenotypic antimicrobial susceptibility data. Submission date was considered instead of collection date when the latter was not available, however, this occurred in only a few cases (<0.5%).

### Creation of the N. gonorrhoeae Pathogenwatch Scientific Steering Group

International experts in the field of *N. gonorrhoeae* AMR, microbiology, genetics, genomics, epidemiology and public health were approached and agreed to participate as members of the ‘*N. gonorrhoeae* Pathogenwatch Scientific Steering Group’ in order to discuss the analytics in Pathogenwatch and make sure they met the current needs of the public health and scientific community. During the updates made to the platform and the preparation of this manuscript, these experts participated in virtual sessions to discuss the list of genetic AMR determinants and their association with SIR categories (Table 2) based on experimental and/or computational evidence. Some of the members of the group had previously been directly involved in many of these studies. Other current and future updates were also discussed, such as the inclusion of the NG-STAR typing scheme (51) and the organization of published genomes into public collections, data sharing, privacy and the interconnectivity of Pathogenwatch with other platforms, such as PubMLST (48) or the ENA. The group will regularly discuss new updates to the platform.

**Table 2.** List of *N. gonorrhoeae* genetic antimicrobial resistance (AMR) determinants in Pathogenwatch. References that report evidence of association of each mechanism to AMR in clinical isolates and/or where their role on AMR has been confirmed in the laboratory through, e.g. transformation experiments, are included in the table. Effect: R = resistance, I = susceptibility but increased exposure, A = additive effect, N = negative effect. R and I follow the EUCAST clinical breakpoints except for azithromycin, for which the epidemiological cut-off (ECOFF) is reported and used instead.

| Antibiotic<br>(MIC breakpoint<br>mg/L) | Genetic AMR determinants | Effect | Evidence<br>(References) |
| --- | --- | --- | --- |
| Azithromycin<br>(R: MIC>1, ECOFF) | 23S rDNA 2045A>G substitution (2059A>G in *E. coli*) | R | (74) |
|  | 23S rDNA 2597C>T substitution (2611C>T in *E. coli*) | R | (90) |
|  | *ermA*, *ermB*, *ermC*, *ermF* genes | R | (91, 92) |
|  | *ereA*, *ereB* genes | R | (22) |
|  | *mefA* gene | R | (92, 93) |
|  | *macAB* promoter -48G>T substitution* | R | (94) |
|  | *mtrR* promoter mosaic** |  |  |
|  | *N. meningitidis*-like mosaic (n=1) | R | (23) |
|  | *N. lactamica*-like mosaic (n=2) | R | (23) |
|  | *mtrD* mosaic** |  |  |
|  | *N. meningitidis*-like mosaic (n=1) | R | (23) |
|  | *N. lactamica*-like mosaic (n=2) | R | (23) |
|  | *mtrR* promoter -57delA* | A | (95, 96) |
|  | *mtrR* G45D | A | (97, 98) |
|  | *mtrC* loss-of-function | N | (26) |
|  | *rpIV* ARAK tandem duplication (position 90) | R | (18) |
|  | *rpIV* KGPSLK tandem duplication (position 83) | R | (18) |
|  | *rpID* G70D | A | (25) |
| Ceftriaxone***<br>(R: MIC>0.125) | *penA* mosaic (A311V, I312M, V316P/T, T483S and G545S) | R | (99-101) |
|  | *penA* V316P, T483S, A501P/V, G542S | R | (99, 100) |
|  | *rpoB* P157L, G158V, R201H | R | (21) |
|  | *rpoD* D92-95 deletion, E98K | I | (21) |
| Cefixime***<br>(R: MIC>0.125) | *mtrR* G45D | A | (97, 98) |
|  | *penA* mosaic (I312M, V316T, G545S) | R | (99-101) |
|  | *penA* mosaic (A311V, I312M, V316P/T, T483S and G545S) | R | (99-101) |
|  | *penA* V316P, T483S, A501P | I | (99, 100) |
|  | *rpoB* P157L, G158V, R201H | I | (21) |
|  | *rpoD* D92-95 deletion, E98K | I | (21) |
| Ciprofloxacin<br>(I: 0.03<MIC≤0.06;<br>R: MIC>0.06) | *gyrA* S91F, D95A/N | R | (102) |
|  | *gyrA* D95G | I | (102) |
|  | *norM* promoter -7A>G, -104C>T substitutions* | I | (103) |
|  | *parC* D86N, S87R | R | (102) |
|  | *parC* S87I/N, S88P, E91K | I | (102) |
|  | *parE* G410V | I | (104) |
| Tetracycline****<br>(I: 0.5<MIC≤1;<br>R: MIC>1) | *mtrR* A39T, G45D | A | (97, 98) |
|  | *mtrR* loss-of-function | I | (22) |
|  | *mtrR* promoter -56A>C substitution, -57delA deletion* | I | (23, 95, 96) |
|  | *mtrR* promoter -131G>A (*mtrC* -120G>A substitution, *mtr120*)* | I | (97) |
|  | *rpsJ* V57M | I | (105) |
|  | *tetM* gene | R | (106) |
| Penicillins<br>(I: 0.06<MIC≤1;<br>R: MIC>1) | *bla*TEM gene | R | (107) |
|  | *mtrR* G45D | I | (97, 98) |
|  | *mtrR* A39T | A | (97) |
|  | *mtrR* loss-of-function | I | (22) |
|  | *mtrR* promoter -56A>C, -57delA* | I | (23, 96) |
|  | *mtrR* promoter -131G>A (*mtrC* -120G>A substitution, *mtr120*)* | I | (97) |
|  | *penA* I312M, V316P/T, ins346D, T483S, A501P/T/V, G542S, G545S, P551S | I | (99, 100) |
|  | *penA* mosaic (I312M, V316T, G545S) | A | (99-101) |
|  | *ponA1* L421P | I | (108) |
|  | *porB1b* G120K, A121N/D | I | (109) |
| Spectinomycin<br>(R: MIC>64) | 16S rDNA 1184C>T (1192C>T in *E. coli*) | R | (110) |
|  | *rpsE* T24P | R | (111) |
|  | *rpsE* V27- deletion, K28E | R/A | (111) |
| Sulfonamides<br>***** | *folP* R228S | R | (22, 112) |

### Data sharing and privacy

Sequencing data and metadata files uploaded to Pathogenwatch by the user are kept within the user’s private account. Genomes can be grouped into collections and these can be toggled between private and accessible to collaborators via a URL. Collection URLs include a 12-letter random string to secure them against brute force searching. Setting a collection to ‘off-line mode’ allows users to work in challenging network conditions, which may be beneficial in LMICs – all data are held within the browser. Users can also integrate private and potentially confidential metadata into the display without uploading it to the Pathogenwatch servers (locally within the browser on a user’s machine).

## Results

### N. gonorrhoeae genome analytics in Pathogenwatch

Pathogenwatch is a web-based platform for epidemiological surveillance using genome sequencing data. After upload, different analytics are run simultaneously (Figure 1): cgMLST (46), MLST (45), NG-MAST (49) and NG-STAR (51) typing schemes (Table 1), a genotypic prediction of phenotypic resistance using a customized AMR library (Table 2) that includes known genetic AMR mechanisms for 8 antimicrobials, as well as statistics on the quality of the assemblies (Additional file 3: Figure S1). These analytical features differentiate Pathogenwatch from a parallel platform from the same group, Microreact (37), which shares one of the main layouts with Pathogenwatch (a phylogenetic tree, a map and a table or timeline), but it is intended for visualization of pre-computed phylogenetic trees with accompanying metadata, while Pathogenwatch also includes analytical tools.

**Figure 1.**
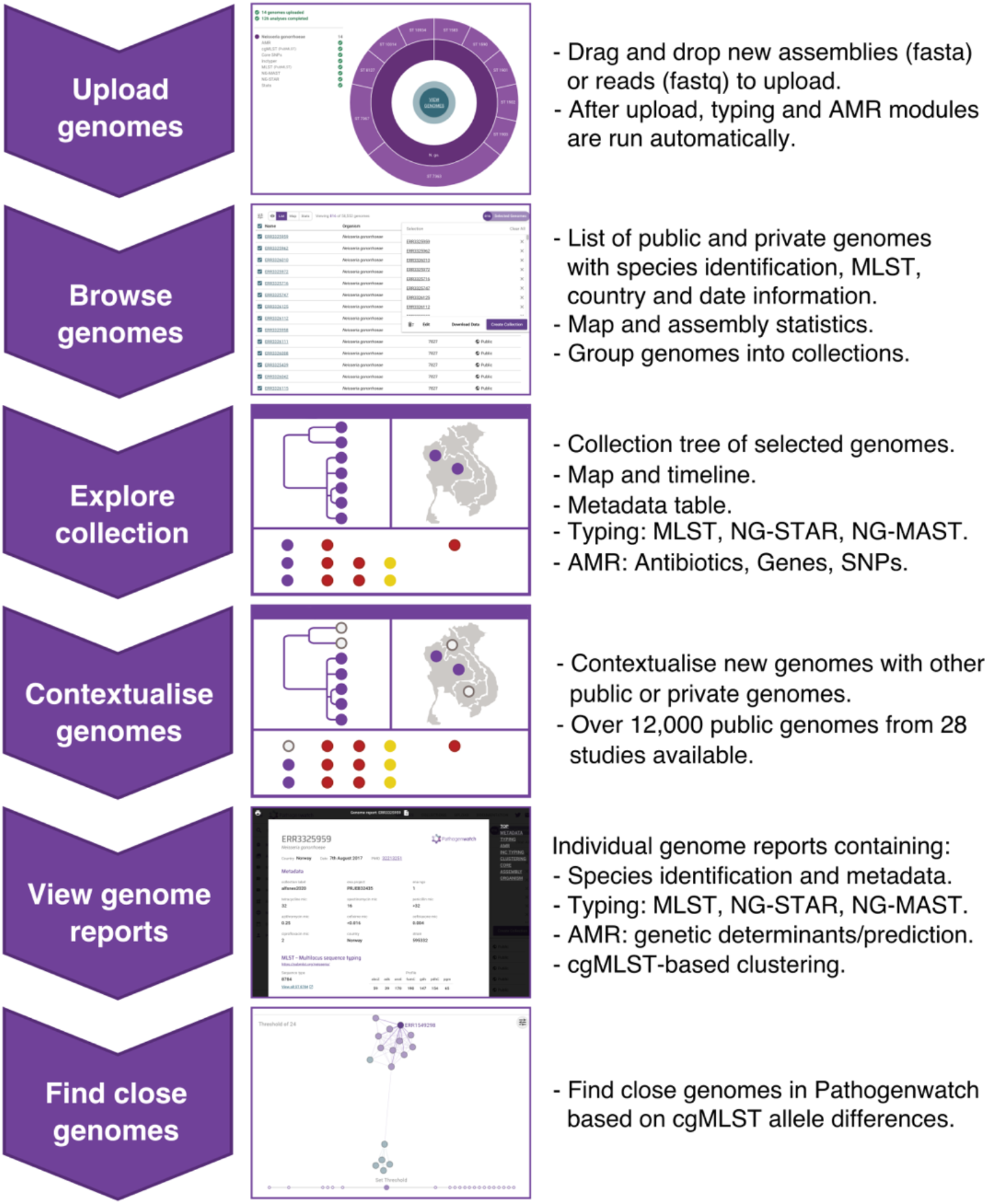
Main workflow in Pathogenwatch. New genomes can be uploaded and combined with public data for contextualisation. The collection view allows data exploration through a combined phylogenetic tree, a map, a timeline and the metadata table, which can be switched to show typing information (Multi-Locus Sequence Typing, MLST; *N. gonorrhoeae* Sequence Typing for Antimicrobial Resistance, NG-STAR; and *N. gonorrhoeae* Multi-Antigen Sequence Typing, NG-MAST) as well as known genetic AMR mechanisms for eight antibiotics. Genome reports summarise the metadata, typing and AMR marker results for individual isolates and allow finding other close genomes in Pathogenwatch based on core genome MLST (cgMLST). SNPs: single nucleotide polymorphisms.

Genomes from one or multiple studies can be grouped into collections (Figure 2 and Additional file 3: Figure S2), and the genomic data are automatically processed by comparing with a core *N. gonorrhoeae* genome built from WHO reference strain genomes (15, 22). A phylogenetic tree is obtained as a result, representing the genetic relationship among the isolates in the collection. Metadata can be uploaded at the same time as the genome data, and if the collection location coordinates for an isolate are provided, this information is plotted into a map (Additional file 3: Figure S1). If date or year of isolation is also provided, this information is represented in a timeline. The three panels on the main collection layout - the tree, the map and a table or timeline – are functionally integrated so filters and selections made by the user update all of them simultaneously. Users can also easily switch among the metadata and the results of the main analytics: typing, genome assembly statistics, genotypic AMR prediction, AMR-associated SNPs, AMR-associated genes and the timeline (Additional file 3: Figure S1). cgMLST is used for finding close genomes in the database based on allele differences to one individual isolate (Additional file 3: Figure S3). A video demonstrating the usage and main features of Pathogenwatch is available (87). Notes on data sharing and privacy are available in the Methods section.

**Figure 2.**
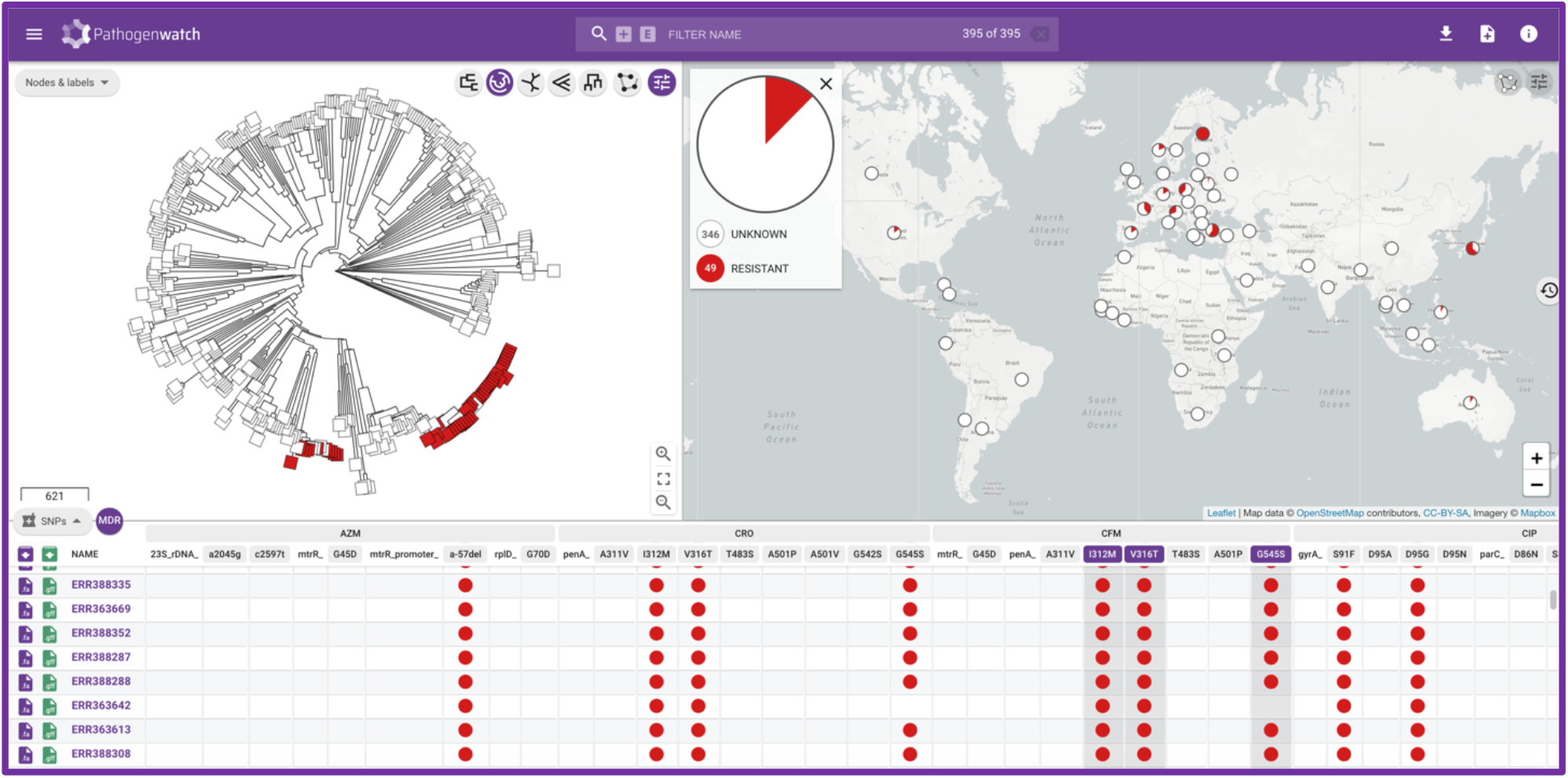
Main display of a Pathogenwatch collection, showing a phylogenetic tree, a map and a table of SNPs associated with AMR of 395 *N. gonorrhoeae* genomes from a global study (64, 88). Isolates carrying three mosaic *penA* marker mutations are marked in red in the tree and the map. The table can be switched to show the metadata, a timeline, typing results (Multi-Locus Sequence Typing, MLST; *N. gonorrhoeae* Sequence Typing for Antimicrobial Resistance, NG-STAR and *N. gonorrhoeae* Multi-Antigen Sequence Typing, NG-MAST) as well as AMR analytics (known genetic mechanisms and genotypic AMR prediction) implemented in the platform. Further detail is shown in Additional file 3: Figure S1.

### Library of genetic AMR mechanisms: genotypic and phenotypic benchmarks

We compiled described genetic AMR mechanisms previously reported for *N. gonorrhoeae* up to the writing of this manuscript into the AMR library in Pathogenwatch (Table 2).A genotypic accuracy testing of the AMR library was performed using the 14 *N. gonorrhoeae* reference genomes from the WHO 2016 panel (22), which were uploaded into Pathogenwatch. All the genetic AMR determinants described as present in these isolates and implemented in the Pathogenwatch AMR library were obtained as a result (Additional file 1: Table S2). Only one discrepancy was found when compared to the original publication. The WHO U strain was reported as carrying a *parC* S87W mutation. However, mapping the original Illumina data from this isolate with the final genome assembly revealed that this strain carries a wild type allele (Additional file 3: Figure S4). MLST and NG-MAST types were the same as those reported in the original publication (note that NG-STAR was not available at that time) and the *porA* mutant gene was found in WHO U as previously described. This mutant *porA* has nearly a 95% nucleotide identity to *N. meningitidis* and 89% to *N. gonorrhoeae*, and it is included as screening because it has previously been shown to cause false negative results in some molecular detection tests for *N. gonorrhoeae* (114).

Then, we also performed a genotypic-phenotypic benchmark using a test dataset of 3,987 *N. gonorrhoeae* isolates from 13 different studies containing MIC information for at least part of the following six antibiotics: ceftriaxone, cefixime, azithromycin, ciprofloxacin, benzylpenicillin and tetracycline (Additional file 1: Table S1). EUCAST clinical breakpoints were applied for five of the antimicrobials except for azithromycin, for which the adoption of an ECOFF>1 mg/L is now recommended to distinguish isolates with azithromycin resistance determinants, instead of a clinical resistance breakpoint (115, 116). A visualization of the range of MICs on each particular combination of genetic AMR mechanisms observed on the isolates from the benchmark test dataset (Figure 3a-b and Additional file 3: Figures S5-S10) revealed combinations that show an additive effect on AMR. These combinations were included in the AMR library to improve the accuracy of the genotypic prediction. For example, *rpsJ* V57M and some *mtrR*-associated mutations individually are associated with a decreased susceptibility or intermediate resistance to tetracycline (MICs of 0.5-1 mg/L), however, a combination of these variants can increase MICs above the EUCAST resistance breakpoint for tetracycline (MICs>1 mg/L) (Additional file 3: Figure S9). This is the case of the combination of *rpsJ* V57M with the *mtrR* promoter -57delA mutation (N=681 isolates, 94.9% positive predictive value, PPV) or with *mtrR* promoter -57delA and *mtrR* G45D (N=83 isolates, 93.9% PPV). Several combinations of *penA, ponA1, mtrR* and *porB1b* mutations were observed to be able to increase the benzylpenicillin MIC above the resistant threshold in most of the cases (Additional file 3: Figure S10). This is the case of the *porB1b* mutations combined with *mtrR* A39T (N=31 isolates, 100% PPV), with the *mtrR* promoter -57delA deletion (N=286 isolates, 96.5% PPV) or with *mtrR* promoter -57delA and *ponA1* L421P (N=269 isolates, 96.3%). Despite mosaic *penA* not being a main driver of resistance to penicillins, a combination of the *porB1b* mutations with the three main mosaic *penA* mutations (G545S, I312M and V316T) was also associated with a resistant phenotype in all cases (N=17 isolates, 100% PPV). A recent publication showed that loss-of-function mutations in *mtrC* increased susceptibility to azithromycin and are associated with isolates from the cervical environment (26). We included the presence of a disrupted *mtrC* as a modifier of antimicrobial susceptibility in the presence of an *mtr* mosaic, as we did not have enough evidence from the test dataset to assess the MIC ranges of isolates with the 23S rDNA A2045G and C2597T mutations with and without a disrupted *mtrC* gene.

**Figure 3.**
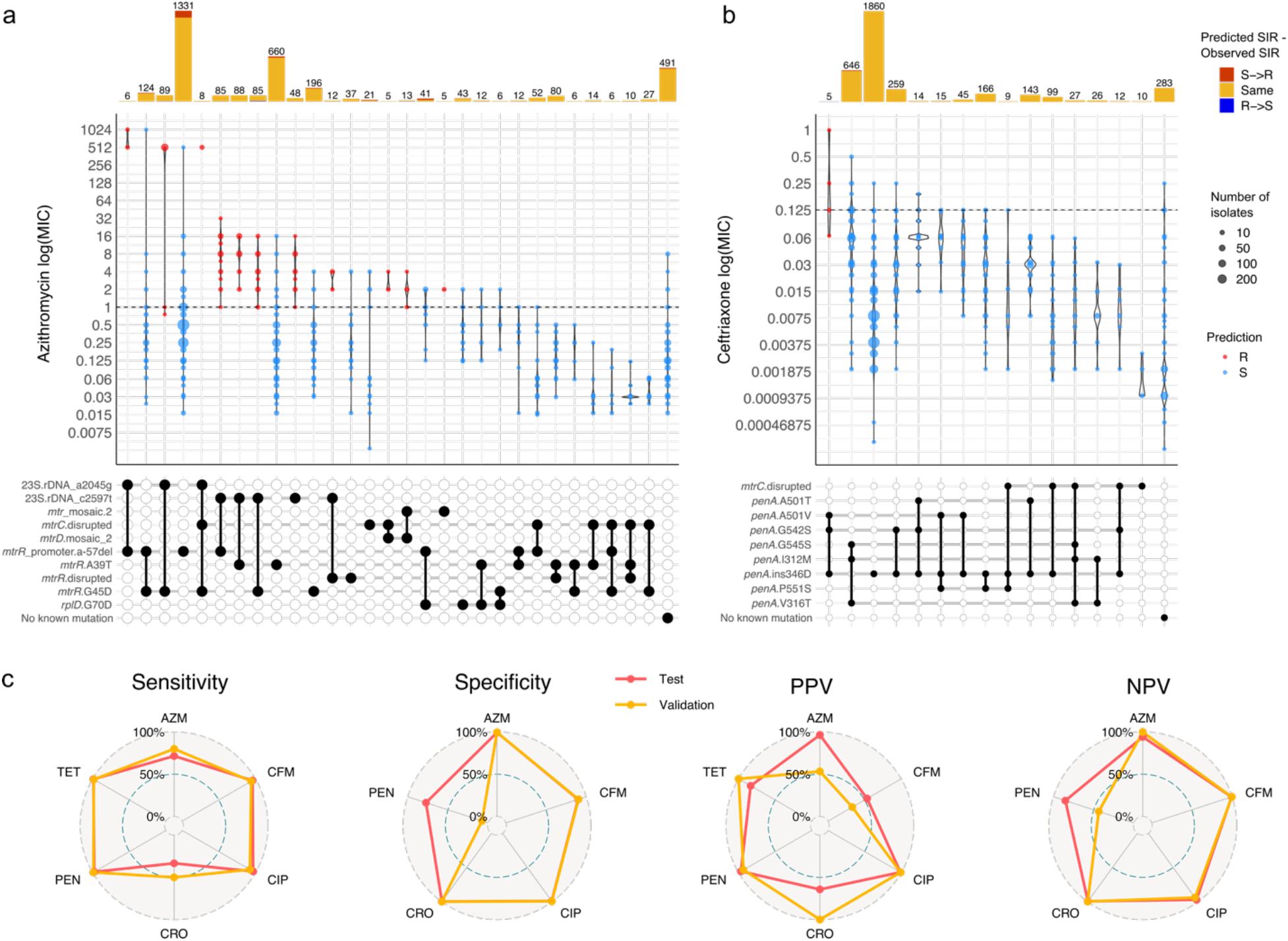
Distribution of minimum inhibitory concentration (MIC) values (mg/L) for the last-line antibiotics for *N. gonorrhoeae* azithromycin (a) and ceftriaxone (b) in a collection of 3,987 *N. gonorrhoeae* isolates with different combinations of genetic antimicrobial resistance (AMR) mechanisms. Only combinations observed in at least 5 isolates are shown (see Additional file 3: Figure S5-S10 for expanded plots for six antibiotics). Dashed horizontal lines on the violin plots mark the EUCAST epidemiological cut-off (ECOFF) for azithromycin and EUCAST clinical breakpoint for ceftriaxone. Point colours inside violins represent the genotypic AMR prediction by Pathogenwatch on each combination of mechanisms (indicated by black circles connected vertically; horizontal thick grey lines connect combinations of mechanisms that share an individual determinant). Barplots on the top show the abundance of isolates with each combination of mechanisms. Bar colours represent the differences between the predicted and the observed SIR (i.e. red for a predicted susceptible mechanism when the observed phenotype is resistant). (c) Radar plots comparing the sensitivity, specificity, positive and negative predictive values (PPV/NPV) for six antibiotics for the test and validation benchmark analyses. AZM = Azithromycin, CFM = Cefixime, CIP = Ciprofloxacin, CRO = Ceftriaxone, PEN = Benzylpenicillin, TET = Tetracycline.

Results from the benchmark (Additional file 1: Table S3) show sensitivity values (true positive rates, TP/(TP+FN); TP=True Positives, FN=False Negatives) above 96% for tetracycline (99.2%), benzylpenicillin (98.1%), ciprofloxacin (97.1%) and cefixime (96.1%), followed by azithromycin (71.6%) and ceftriaxone (33.3%). These results reflect the complexity of the resistance mechanisms for azithromycin and ceftriaxone, where the known genetic determinants explain only part of the antimicrobial susceptibility. However, specificity values (true negative rates, TN/(TN+FP); TN=True Negatives, FP=False Positives) for these two antibiotics as well as ciprofloxacin were above 99% (Additional file 1: Table S3), demonstrating that the genetic mechanisms included in the database have a role in AMR. The specificity value for cefixime was lower but nearly 90%, mainly due to the high number of isolates with an MIC below the threshold but with three mutations characterising a mosaic *penA* allele (G545S, I312M and V316T, TP=367, TN=323, PPV=53.2%; Additional file 1: Table S4). Benzylpenicillin and tetracycline showed specificity values of 77.3% and 61.3%, respectively. In the first case, all the mechanisms included in the library showed a PPV value above 94%. For tetracycline, a considerable number of false positive results are mainly caused by the presence of *rpsJ* V57M, for which PPV=83.8% (TP=1083, FP=209; Additional file 1: Table S4). However, this mutation was kept in the AMR library because it can cause intermediate resistance to tetracycline on its own (Additional file 3: Figure S9).

Results from the benchmark analysis on the 3,987-isolates dataset were used to curate and optimize the AMR library. Thus, in order to objectively validate it, the benchmark analysis was also run on a combination of three different collections (N=1,607, Additional file 1: Table S1) with available MIC information for seven antibiotics including spectinomycin (Additional file 1: Table S3) (69, 70, 117). Results from the test and validation benchmark runs were compared, showing that sensitivity values on the six overlapping antibiotics were very similar, with the validation benchmark performing even better for azithromycin and ceftriaxone (Figure 3c). In terms of specificity, both datasets performed equally well for all antibiotics except for benzylpenicillin, in which specificity drops in the validation benchmark. This is due to the *penA*_ins346D mutation (TP=1125, FP=83) and the *blaTEM* genes (TP=525, FP=36), which despite showing false positives, have a PPV above 93% (Additional file 1: Table S5). In general, discrepancies found between the test and the validation benchmarks can be explained by particular mechanisms that on their own show high predictive values and affect antibiotics for which we do not currently understand all the factors involved in resistance, such as azithromycin and the ESCs (Additional file 1: Table S5).

### Over 12,000 public genomes available

Data for 11,461 isolates were successfully assembled and passed all quality cut-offs, resulting in 12,515 isolates after including the previously-available Euro-GASP 2013 dataset (15). New assemblies were uploaded and made public on Pathogenwatch, which now constitutes the largest repository of curated *N. gonorrhoeae* genomic data with associated metadata, typing and AMR information at the time of submission of this manuscript. Updated data spans 27 different publications (18, 53, 58-61, 63-65, 67-70, 117-131) and is organized into individual collections associated with the different studies (Additional file 1: Table S6). Available metadata was added for the genomes from these publications while basic metadata fields were kept for others (country, year/date and ENA project number).

We cross-checked that the main clusters found in the phylogenetic trees obtained after creating the public collections in Pathogenwatch were consistent with those observed in the trees in the corresponding publications. For example, recent works defined two major clusters of *N. gonorrhoeae*, termed Lineages A and B, which were found to be consistent with the corresponding Pathogenwatch trees as exemplified for isolates from England in Town *et al* (2020) (68) (Figure S11a). We were also able to differentiate the cefixime-resistant *penA10* and *penA34*-carrying clones from Vietnam from Lan *et al* (2020) (124) (Figure S11b) as well as the 10 major clusters defined in the *N. gonorrhoeae* population circulating in New York City (NYC) as described in Mortimer *et al* (2020) (120) (Figure S11c). In the last case, we also liked to emphasize the usefulness of Microreact (37) as a parallel tool to Pathogenwatch for more complex visualization purposes, such as showing the 10 major clusters in NYC as metadata blocks of different colours.

The *N. gonorrhoeae* public data available on Pathogenwatch spans nearly a century (1928-2018) and almost 70 different countries (Additional file 3: Figure S12). However, sequencing efforts are unevenly distributed around the world, and over 90% of the published isolates were isolated in only 10 countries, headed by the United Kingdom (N=3,476), the United States (N=2,774) and Australia (N=2,388) (Additional file 1: Table S7, Figure 4). A total of 554 MLST, 1,670 NG-MAST and 1,769 NG-STAR different STs were found in the whole dataset, from which a considerable number were new profiles caused by previously undetected alleles or new combinations of known alleles (N=92 new MLST STs, N=769 new NG-STAR STs and N=2,289 isolates with new NG-MAST *porB* and/or *tbpB* alleles). These new alleles and profiles were submitted to the corresponding scheme servers.

**Figure 4.**
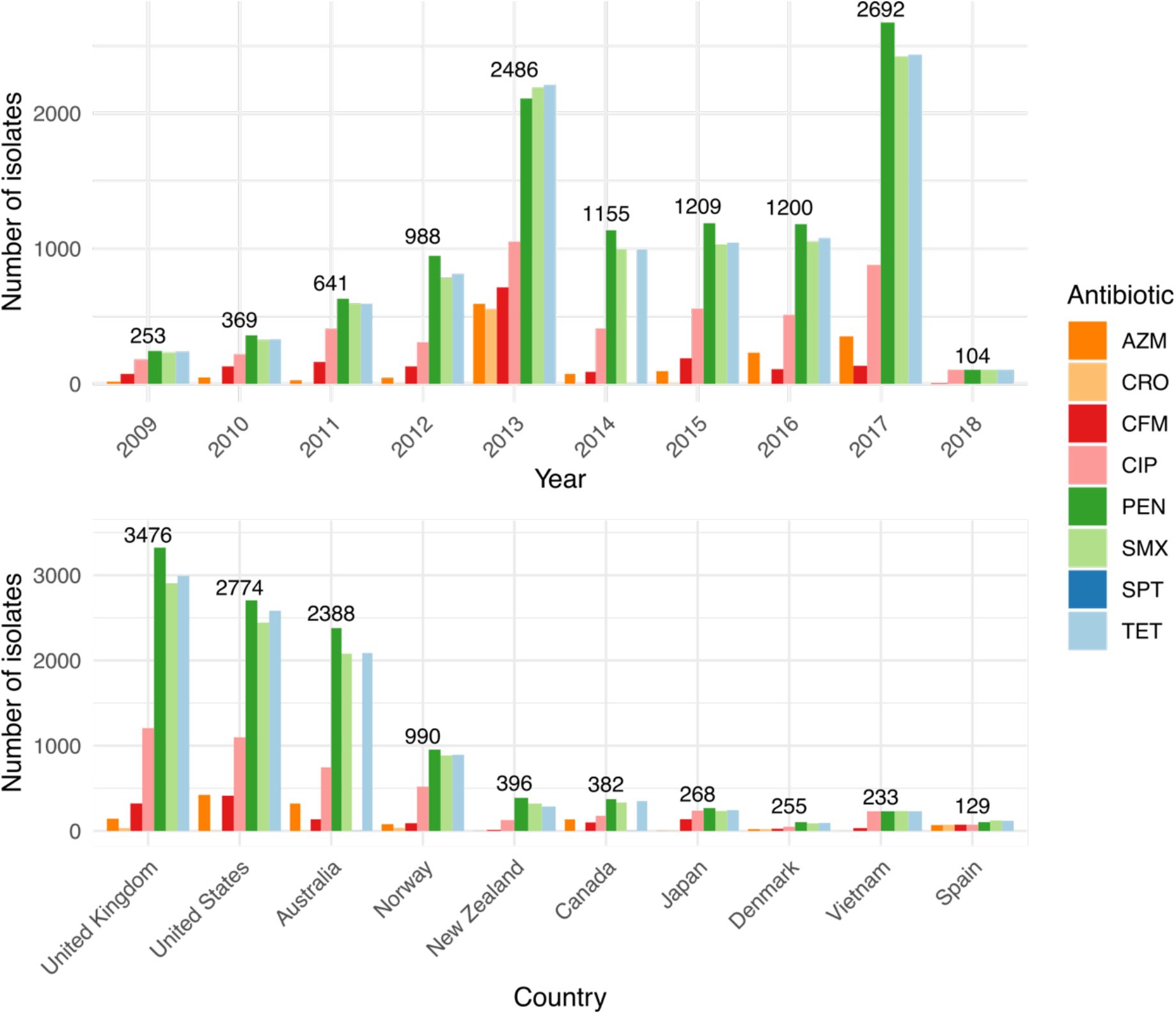
Summary of the geolocalization and collection date of 12,515 public *N. gonorrhoeae* genomes in Pathogenwatch. Coloured bars represent the genotypic antimicrobial resistance (AMR) prediction based on the mechanisms included in the library. AZM = Azithromycin, CFM = Cefixime, CIP = Ciprofloxacin, CRO = Ceftriaxone, PEN = Benzylpenicillin, TET = Tetracycline.

Genomic studies are often biased towards AMR isolates, and this is reflected in the most abundant STs found for the three typing schemes within the public data. Isolates with MLST ST1901, ST9363 and ST7363, which contain resistance mechanisms to almost every antibiotic included in the study, represent over 25% of the data (Figure 5). Isolates with MLST ST1901 and ST7363 are almost always associated with resistance to tetracycline, sulfonamides, benzylpenicillin and ciprofloxacin and nearly 50% of isolates from these two types harbour resistance mechanisms to cefixime. Ciprofloxacin resistance is not widespread among ST9363 isolates, which are associated with azithromycin resistance in nearly 50% of the isolates for this ST (Figure 5). NG-STAR ST63 (carrying the non-mosaic *penA-2* allele, *penA* A517G and *mtrR* A39T mutations as described in (52)) is the most represented in the dataset and carries resistance mechanisms to tetracycline, sulfonamides, and benzylpenicillin, but is largely susceptible to spectinomycin, ciprofloxacin, the ESCs cefixime and ceftriaxone and azithromycin. NG-STAR ST90 isolates, conversely, are largely associated with resistance to cefixime, ciprofloxacin and benzylpenicillin as they carry the key resistance mutations in mosaic *penA-34*, as well as in the *mtrR* promoter, *porB1b, ponA, gyrA* and *parC* (as described in (52)). NG-MAST ST1407 is commonly associated with MLST ST1901 and is the second most represented ST in the dataset following NG-MAST ST2992, which mainly harbours resistance to tetracycline, benzylpenicillin and sulfonamides (Figure 5).

**Figure 5.**
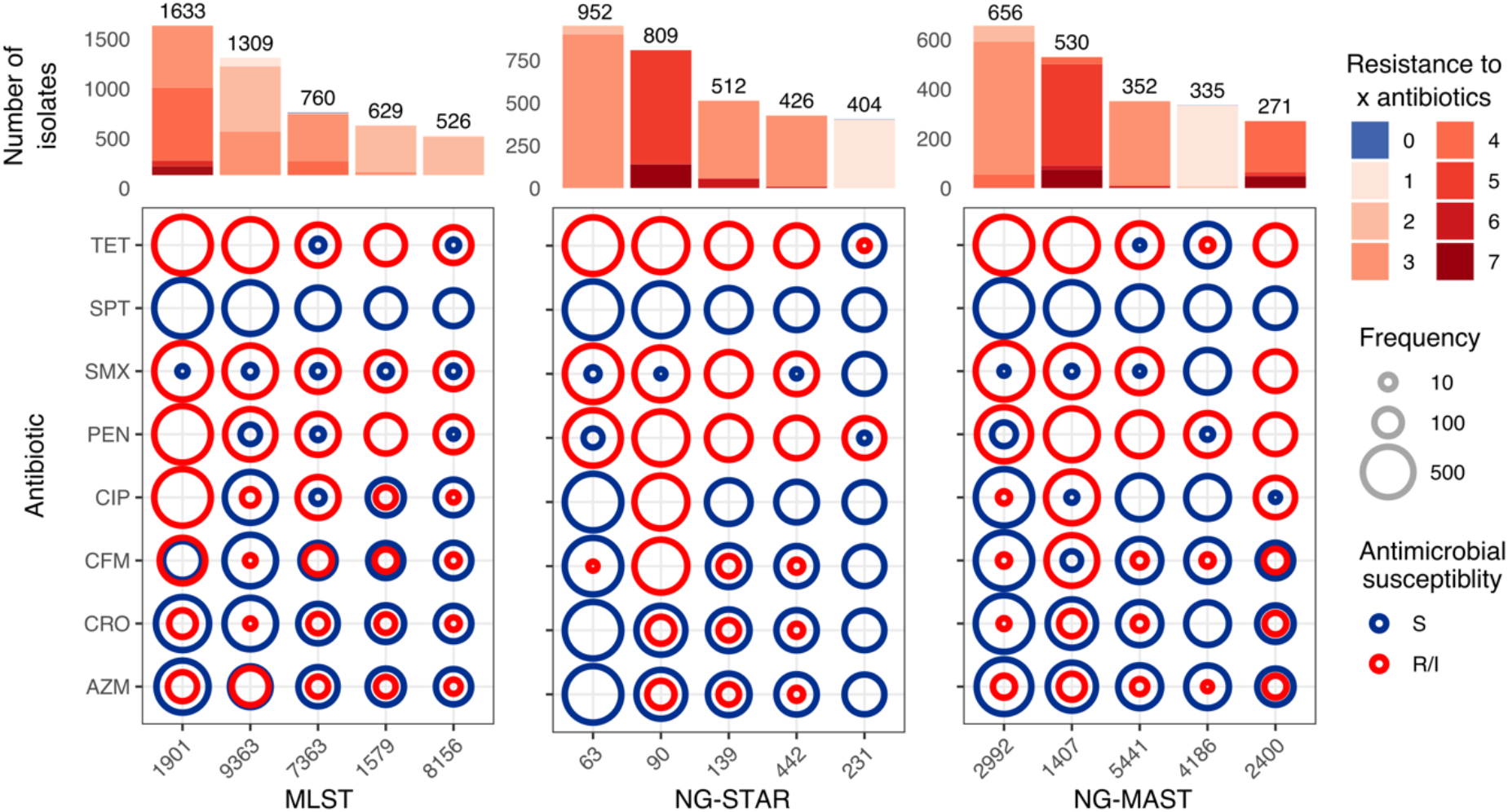
Predicted antimicrobial resistance (AMR) profiles of the top five Multi-Locus Sequence Typing (MLST), *N. gonorrhoeae* Sequence Typing for Antimicrobial Resistance (NG-STAR) and *N. gonorrhoeae* Multi-Antigen Sequence Typing (NG-MAST) types in the *N. gonorrhoeae* public data in Pathogenwatch. Coloured circles in the grids show the proportion of genomes from each ST which are predicted to have an intermediate (susceptible but increased exposure) or resistant phenotype, red) versus susceptible genomes (in dark blue) from each sequence type (ST) and antibiotic. Bars on the top show the number of isolates from each ST coloured by the number of antibiotics the genomes are predicted to be resistant to.

### Case study: global expansion of an mtr mosaic-carrying clone

The genetic mechanisms that have commonly been associated with an increased MIC of azithromycin in *N. gonorrhoeae* are two mutations in the 23S rRNA gene (A2045G and C2597T substitutions, in *N. gonorrhoeae* nomenclature) as well as mutations in *mtrR* and its promoter (132, 133). As described above, other mechanisms have also been recently discovered that increase the MIC of azithromycin (Table 2), such as mosaicism affecting the efflux pump-encoding *mtrCDE* genes and its repressor *mtrR*, mainly when the mosaic spans the *mtrR* promoter region and *mtrD* gene (23, 24). Some studies have recently reported the local expansion of azithromycin-resistant *N. gonorrhoeae* lineages carrying an *mtr* mosaic in the USA (122, 123, 134) and Australia (118). However, the extent of the dispersion of this mechanism to other parts of the world has not been studied yet. Here, using the public genomes of *N. gonorrhoeae* in Pathogenwatch, we have been able to explore this question.

A total of 1,142 strains with genetic determinants of azithromycin resistance were selected in Pathogenwatch and combined with 395 genomes from a global collection (64) for background contextualization (see Pathogenwatch project in (135)) (Figure 6a). 571 of the strains predicted to be resistant to azithromycin had some form of mosaic in the *mtrR* promoter and/or *mtrD* gene of one of the three types described in Wadsworth *et al*. (2018) (23) and included in the Pathogenwatch AMR library (Table 2). These mosaics have been experimentally proven to increase MIC of azithromycin above 1 mg/L, which is the EUCAST ECOFF value as well as the Clinical Laboratory and Standards Institute (CLSI) non-susceptibility breakpoint (23, 24). One of the *N. lactamica*-like mosaics, termed here ‘*mtr*_mosaic.2’, was by far the most extended, as it was found in 545 genomes spanning the *mtrR* promoter and/or the *mtrD* gene, with 521 (95.6%) of them spanning both regions. Twenty-five genomes contained a *N. meningitidis-*like mosaic *mtrR* promoter and/or *mtrD* gene (‘*mtr_*mosaic.1’) and in only 9 (36%) of them the mosaic spanned both loci. The *N. lactamica*-like ‘*mtr_*mosaic.3’ was only found in isolate ERR855360 (GCGS834) from Los Angeles (USA, 2012), which is where the reference sequence for this mosaic was extracted from. Of the studies where these *mtr* mosaic-carrying genomes were obtained from, only those from the USA and Australia specifically targeted and found this genetic determinant of resistance. The rest did not target this mosaic and some of them found strains with unexplained increased MICs of azithromycin (69, 121, 129), which could partly be explained by the presence of these *mtr* mosaics.

**Figure 6.**
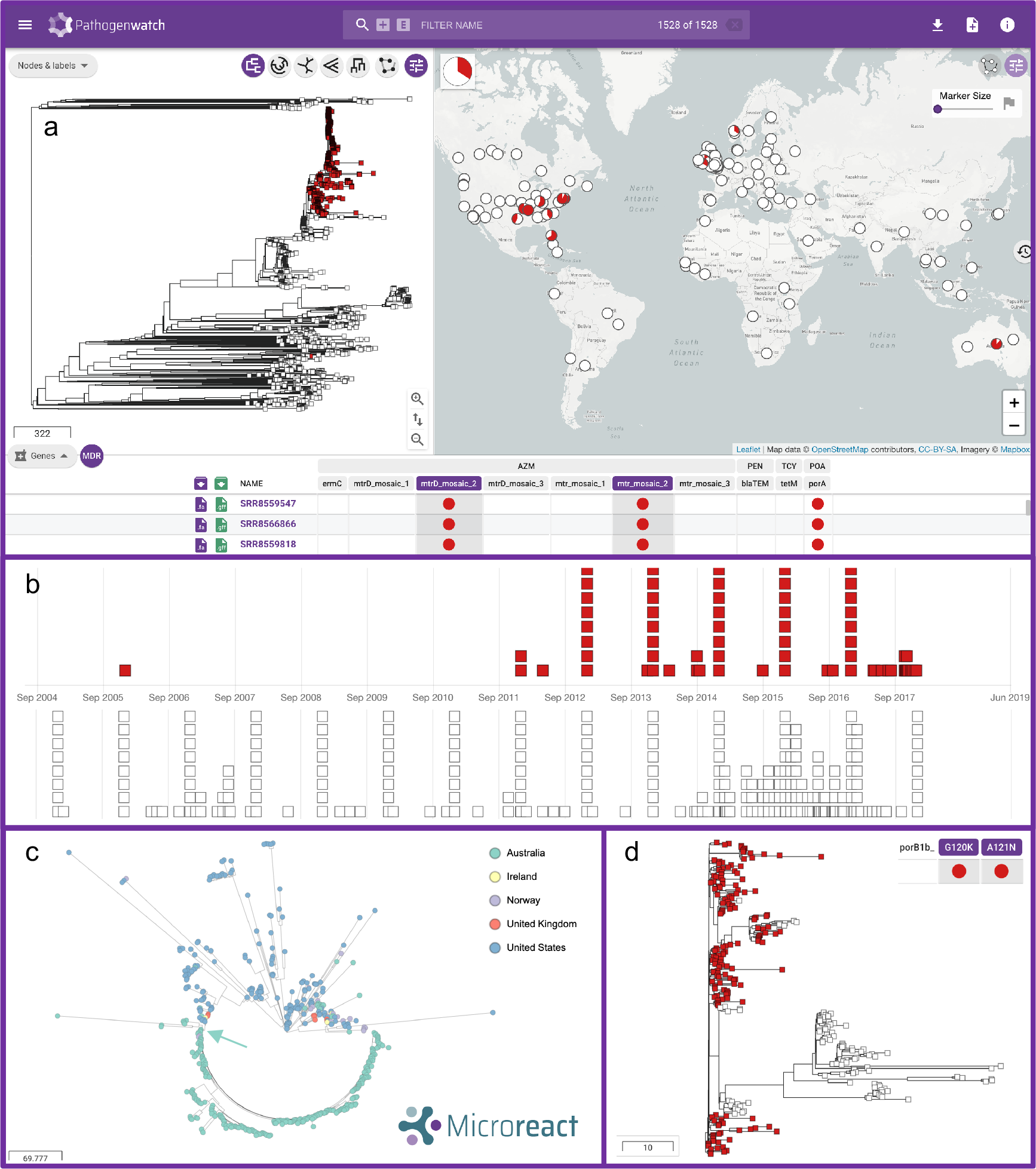
*N. gonorrhoeae* genomes carrying genetic AMR mechanisms associated to azithromycin resistance were selected in Pathogenwatch (n=1,142) and combined with genomes from a global collection (64, 88) (total n=1528) for background contextualization. (a) Main layout of the combined collection, with the emerging lineage carrying *mtr* mosaic 2 spanning the *mtrR* promoter and *mtrD* marked in red in the tree and the map. (b) Timeline of the genomes carrying *mtr* mosaic 2 (in red) and other public genomes in the database without this genetic AMR mechanism. (c) Visualization of the *mtr* mosaic 2-carrying lineage (n=520) spreading in the USA and Australia (see legend) using Microreact. The arrow in turquoise colour marks the divergence of the Australian lineage, shown in more detail in (d) coloured by the presence (in red) or absence (in white) of the *porB1b* G120K and A121N mutations. The Pathogenwatch project of this case study can be explored in (135).

We observed one main lineage carrying mosaic 2 in *mtrR* promoter and *mtrD* gene (Figure 6a) with 520 genomes. Of those, only 3 and 8 isolates carried the 23S rDNA A2045G and C2597T mutations, respectively. Interestingly, the first strain in the database with this type of mosaic dates from 2006 (18), however, it was not until the end of 2011-2012 when this lineage started to expand (Figure 6b). Despite the genomic data contained in Pathogenwatch being biased to the amount of data sequenced and published from each country and year, we can easily infer that this lineage has spread across the world as we detect cases in Australia (n=293) (118), the USA (n=195) (18, 120, 122, 123), Norway (n=19) (121), the United Kingdom (n=11) (68, 119), and Ireland (n=3) (129). A strong association was found to the country of isolation (Figure 6c), with a broad diversity of sublineages having spread across the USA (strains mostly isolated between 2012 and 2016). In contrast, an expansion of a particular clone, likely from a single main introduction, was observed to have occurred in Australia (strains isolated in 2017), followed by a further divergence of a subclone within the country which correlates with the loss of the *porB1b* G120K and A121N mutations (Figure 6d), likely through a recombination event. Despite epidemiological data not being available for the Australian study (118), from their work we know that the clusters carrying an *mtr* mosaic were mostly linked to transmission between men, although bridging among MSM and heterosexual populations was also observed.

The results from our case study show that there is an emerging lineage of *N. gonorrhoeae* that has spread across the world and that is carrying a mosaic *mtr* that has been associated with low-to-medium resistance to azithromycin. This global lineage, as well as others that may emerge carrying this or other genetic AMR mechanisms, has to be closely monitored. For this purpose, an up-to-date genomic epidemiology tool such as Pathogenwatch, which includes a list of genetic AMR mechanisms approved by an expert group is a great resource for the scientific community. At the moment, Pathogenwatch includes references for three types of mosaics in the *mtrR* promoter and *mtrD* genes that have been experimentally proven to increase MIC of azithromycin (23, 24), and the detection of these mosaics on new genomes respond to a set of similarity rules (see Data availability section). However, we will keep the database updated with new experimentally-confirmed reference sequences that may arise from further studies as it is still unclear whether all mosaics affecting the *mtrCDE* efflux pump will cause a decreased susceptibility to azithromycin.

## Discussion

We present a public health focussed *N. gonorrhoeae* framework at Pathogenwatch, an open access platform for genomic surveillance supported by an expert group that can be adapted to any public health or microbiology laboratory. Little bioinformatics expertise is required, and users can choose to either upload raw short read data or assembled genomes. In both cases, the upload of high-quality data is encouraged in the form of quality-checked reads and/or quality-checked assemblies. Recent benchmark analyses show particular recommendations for long-read or hybrid data (136) as well as short read-only data (40, 137). On upload, several analyses are run on the genomes, and results for the three main typing schemes (MLST, NG-MAST and NG-STAR) as well as the detection of genetic determinants of AMR and a prediction of phenotypic resistance using these mechanisms can be obtained simultaneously. The library of AMR determinants contained in Pathogenwatch for *N. gonorrhoeae* has been revised and extended to include the latest mechanisms and epistatic interactions with experimental evidence of decreasing susceptibility or increasing resistance to at least one of eight antibiotics (Tables 2). A test and validation benchmark analyses revealed sensitivity and/or specificity values >90% for most of the tested antibiotics (Additional file 1: Table S3). Sensitivity values for the antimicrobials in the current dual treatment, azithromycin (80%) and ceftriaxone (50%), reflect the complexity of the resistance mechanisms for these antibiotics, for which we can only explain part of the observed phenotypic resistance. However, their specificity values were above 99% (Additional file 1: Table S3), further strengthening the associations of the included AMR determinants in increasing MICs of these antibiotics. It remains essential to perform phenotypic susceptibility testing so we can detect inconsistencies between phenotypic and genotypic data that can lead to the identification and subsequent verification of novel or unknown resistance mechanisms. This will allow to continuously expand the list of genetic AMR mechanisms, and the AMR prediction from genomic data will further improve.

The continuous increase in reporting of *N. gonorrhoeae* AMR isolates worldwide led to a call for international collaborative action in 2017 to join efforts towards a global surveillance scheme. This was part of the WHO global health sector strategy on STIs (2016-2021), which set the goal of ending STI epidemics as a public health concern by year 2030 (7, 8). Several programmes are currently in place at different global, regional or national levels to monitor gonococcal AMR trends, emerging resistances and refine treatment guidelines and public health policies. This is the case of, for example, the WHO Global Gonococcal Antimicrobial Surveillance Programme (WHO GASP) (7, 8), the Euro-GASP in Europe (6, 15, 138), the Gonococcal Isolate Surveillance Project (GISP) in the United States (139), the Canadian Gonococcal Antimicrobial Surveillance Programme (140), the Gonococcal Surveillance Programme (AGSP) in Australia (141) or the Gonococcal Resistance to Antimicrobials Surveillance Programme (GRASP) in England and Wales (142). The WHO in collaboration with CDC has recently started an enhanced GASP (EGASP) (143) in some sentinel countries such as the Philippines and Thailand (144), aimed at collecting standardized and quality-assured epidemiological, clinical, microbiological and AMR data. On top of these programs, WHO launched the Global AMR Surveillance System (GLASS) in 2015 to foster national surveillance systems and enable standardized, comparable and validated AMR data on priority human bacterial pathogens (145). Efforts are now underway to link WHO GASP to GLASS. However, gonococcal AMR surveillance is still suboptimal or even lacking in many locations, especially in LMICs, such as several parts of Asia, Central and Latin America, Eastern Europe and Africa, which worryingly have the greatest incidence of gonorrhoea (3). LMICs often have access to antimicrobials without prescription, have limited access to an optimal treatment, lack the capacity needed to perform a laboratory diagnosis due to limited or non-existent quality-assured laboratories, microbiological and bioinformatics expertise or training, insufficient availability and exorbitant prices of some reagents on top of a lack of funding, which altogether compromises infection control.

High throughput sequencing approaches have proved invaluable over traditional molecular methods to identify AMR clones of bacterial pathogens, outbreaks, transmission networks and national and international spread among others (28, 29). Genomic surveillance efforts to capture the local and international spread of *N. gonorrhoeae* have resulted in several publications within the last decade involving high throughput sequence data of thousands of isolates from many locations across the world. The analysis of this data requires expertise, not always completely available, in bioinformatics, genomics, genetics, AMR, phylogenetics, epidemiology, etc. For lower-resourced settings, initiatives such as the NIHR Global Health Research Unit, Genomic Surveillance of Antimicrobial Resistance (146) are essential to build genomic surveillance capacity and provide the necessary microbiology and bioinformatics training for quality-assured genomic surveillance of AMR.

One of the strengths of genomic epidemiology is being able to compare new genomes with existing data from a broader geographical level, which provides additional information on, e.g. if new cases are part of a single clonal expansion or multiple introductions from outside a specific location. To support this, Pathogenwatch calculates phylogenetic trees from a set of genomes selected as collections. Currently, over 12,000 isolates of *N. gonorrhoeae* have been sequenced using high throughput approaches and publicly deposited on the ENA linked to a scientific publication. We have quality-checked and assembled these data using a common pipeline and we made it available through Pathogenwatch, with the aim of representing as much genomic diversity of this pathogen as possible to serve as background for new analyses. These public genomes are associated with at least 27 different scientific publications, and have been organized in Pathogenwatch as individual collections (Additional file 1: Table S6). The clustering of strains on the resulting reconstructions was found consistent with those in the original publications (some examples in Figure S11), while differences in branch lengths may be attributed to the usage of different reconstruction methods.

The power of Pathogenwatch to investigate questions of public health concern is reflected in a case study (Figure 6). By selecting 1,142 azithromycin resistant strains from the public data in Pathogenwatch in the context of a global collection (64), we observed one clone carrying *N. lactamica*-like *mtr* mosaic (‘mosaic_2’) in both the *mtrR* promoter and *mtrD* genes, likely resulting from the same recombination event. Strong geographical structure was found in these azithromycin resistant strains, with isolates from the USA (mostly from 2012-2016) clearly differentiated from those from Australia (from 2017), which show a more clonal dispersion, likely from a single main introduction to the country followed by a rapid spread. Interestingly, a sublineage of this Australian *mtr* mosaic-carrying clone seems to have also diverged after losing the *porB1b* G120K and D121N mutations. It is important to note that the data from which these inferences were derived was gathered from surveillance-based studies and outbreak investigations, which may bias the observed global diversity of strains carrying this mosaic. Phenotypic susceptibility data for azithromycin or epidemiological information were not available for over half of these strains, thus impeding making further inferences. This reflects the need of improving the submission of anonymized epidemiological and antimicrobial susceptibility data for individual isolates rather than aggregated data to public repositories and/or as supplementary information of the corresponding publications, as this is where the public data in Pathogenwatch is coming from.

In this study, we have additionally gathered an advisory group of *N. gonorrhoeae* experts in different fields such as AMR, microbiology, genetics, genomics, epidemiology and public health who will consult and discuss current and future analytics to be included to address the global public health needs of the community. We suggest this strategy as a role model for other pathogens in this and other genomic surveillance platforms, so the end user, who may not have full computational experience in some cases, can be confident that the analytics and databases underlying this tool are appropriate, and can have access to all the results provided by Pathogenwatch through uploading the data via a web browser. We are aware that this is a constantly moving field and analytics will be expanded and updated in the future. These updates will be discussed within an advisory group to make sure they are useful in the field and the way results are reported is of use to different profiles (microbiologists, epidemiologists, public health professionals, etc.). Future analytics that are under discussion include the automatic submission of new MLST, NG-STAR and NG-MAST STs and alleles to the corresponding servers, e.g. PubMLST (48) and the automatic submission of data to public archives such as the ENA. Inter-connectivity and comparability of results with PubMLST is of particular interest, as this database has traditionally been the reference for *Neisseria* sequence typing and genomics and it is widely used by the *N. gonorrhoeae* community. Plasmid and *tetM/blaTEM* subtyping as recently described (147) will also be considered within the development roadmap of Pathogenwatch. Including a separate library to automatically screen targets of potential interest for vaccine design (148-150) as well as targets of new antibiotics currently in phase III clinical trials (i.e. zoliflodacin (151) or gepotidacin (152)) can also be an interesting addition to the scheme. Regarding AMR, new methods for phenotypic prediction using genetic data are continuously being reported (62, 153, 154), especially those based on machine learning algorithms (155), and will be considered for future versions of the platform. The prediction of MIC values or ranges instead of SIR categories will allow users to decide whether to use EUCAST (156) or CLSI (157) guidelines for categorization.

## Conclusions

In summary, we present a genomic surveillance platform adapted to *N. gonorrhoeae*, one of the main public health priorities compromising the control of AMR infections, where decisions on existing and updated databases and analytics as well as how results are reported will be discussed with an advisory board of experts in different public health areas. This will allow scientists from both higher or lower resourced settings with different capacities regarding high throughput sequencing, bioinformatics and data interpretation, to be able to use a reproducible and quality-assured platform where analyse and contextualise genomic data resulting from the investigation of treatment failures, outbreaks, transmission chains and networks at different regional scales. This open access and reproducible platform constitutes one step further into an international collaborative effort where countries can keep ownership of their data in line with national STI and AMR surveillance and control programs while aligning with global strategies for a joint action towards battling AMR *N. gonorrhoeae*.

## List of abbreviations

AGSP: Australian Gonococcal Surveillance Programme
AMR: Antimicrobial Resistance
AZM: Azithromycin
CDC: Centers for Disease Control and Prevention
CFM: Cefixime
cgMLST: Core Genome Multi-Locus Sequence Typing
CIP: Ciprofloxacin
CLSI: Clinical Laboratory and Standards Institute
CRO: Ceftriaxone
ECOFF: Epidemiological Cut-Off
EGASP: Enhanced Gonococcal Antimicrobial Surveillance Programme
ENA: European Nucleotide Archive
ESCs: Extended Spectrum Cephalosporins
EUCAST: European Committee on Antimicrobial Susceptibility Testing
Euro-GASP: European Gonococcal Antimicrobial Surveillance Programme
FN: False Negative
FP: False Positive
GASP: Gonococcal Antimicrobial Surveillance Programme
GISP: Gonococcal Isolate Surveillance Project
GRASP: Gonococcal Resistance to Antimicrobials Surveillance Programme
HIV: Human Immunodeficiency Virus
LMICs: Low and Middle-Income Countries
MIC: Minimum Inhibitory Concentration
MLST: Multi-Locus Sequence Typing
NG-MAST: *N. gonorrhoeae* Multi-Antigen Sequence Typing
NG-STAR: *N. gonorrhoeae* Sequence Typing for Antimicrobial Resistance
NPV: Negative Predictive Value
PEN: Benzylpenicillin
PPV: Positive Predictive Value
SNPs: Single Nucleotide Polymorphisms
ST: Sequence Type
STI: Sexually-Transmitted Infection
TET: Tetracycline
TN: True Negative
TP: True Positive
UK: United Kingdom
WGS: Whole Genome Sequencing
WHO: World Health Organization

## Declarations

### Ethics approval and consent to participate

Not applicable.

### Consent for publication

Not applicable.

### Availability of data and materials

The assemblies included in the current version of the *N. gonorrhoeae* Pathogenwatch scheme and used for the AMR benchmark analyses were generated from raw sequencing data stored in the ENA. Project accession numbers are included in Additional File 1: Tables S1 and S6. The generated assemblies can be downloaded from Pathogenwatch. The AMR library can be accessed from: https://gitlab.com/cgps/pathogenwatch/amr-libraries/-/blob/master/485.toml. The code to reproduce the figures and analyses in this manuscript can be found in https://gitlab.com/cgps/pathogenwatch/publications/-/tree/master/ngonorrhoeae.

## Authors’ contributions

DMA conceived the Pathogenwatch application. CY, RG, KA, BT, AU and DMA developed the Pathogenwatch application. LSB and DMA contributed to the conception and design of the work. CY and LSB generated, updated and benchmarked the *N. gonorrhoeae* AMR library. BT, CY, AU and LSB obtained, quality-checked and reassembled the raw data from the ENA. LSB revised the assembled data, obtained all metadata available from the corresponding scientific publications and created collections. LSB, CY and DMA analysed the data. LSB and DMA drafted the manuscript. LSB, DMA, CY, SA, KCM, TDM, DG, MJC, YHG, IM, BHR, WMS, GS, KT, TW, SRH and MU contributed to the acquisition, technical and scientific interpretation and discussion of the data. LSB, DMA, MJC, YHG, IM, BHR, WMS, GS, KT, TW and MU agreed to participate in the *N. gonorrhoeae* Pathogenwatch Scientific Steering Group before the preparation of this manuscript, and participated in virtual discussions. All authors read and approved the final manuscript.

### Acknowledgements

We would like to thank MJC, YHG, IM, BHR, WMS, GS, KT, TW and MU for their support on the development of the *N. gonorrhoeae* Pathogenwatch scheme and the creation of the *N. gonorrhoeae* Pathogenwatch Scientific Steering Group.

