## Additional file 1 - Supplementary Tables for "A community-driven resource for genomic epidemiology and antimicrobial resistance prediction of *Neisseria gonorrhoeae* at Pathogenwatch"

Table S1. List of studies included in the antimicrobial resistance (AMR) benchmark analyses. The number of isolates included from each study and the antibiotics for which minimum inhibitory concentration (MIC) information is provided is also indicated.

| Benchmark (N) | Study (Reference) | Number of isolates | Project accession | Antibiotics |
|---|---|---|---|---|
| Test (N=3,987) | Chisholm *et al.* 2015 (58) | 15 | PRJEB14933 | Azithromycin, Ceftriaxone |
| | Golparian *et al.* 2020 (59) | 183 | PRJEB4024 | Azithromycin, Ceftriaxone, Cefixime, Ciprofloxacin, Tetracycline, Benzylpenicillin |
| | Demczuk *et al*. 2015, 2016 (60, 61) | 382 | PRJNA298332 PRJNA266539 | Azithromycin, Ceftriaxone, Cefixime, Ciprofloxacin, Tetracycline, Benzylpenicillin |
| | Harris *et al*. 2018 (15) | 1,054 | PRJEB9227 | Azithromycin, Ceftriaxone, Cefixime, Ciprofloxacin |
| | Eyre *et al*. 2017 (62) | 249 | PRJNA315363 | Azithromycin, Cefixime, Ciprofloxacin, Tetracycline, Benzylpenicillin |
| | Fifer *et al*. 2018 (63) | 101 | PRJEB23008 | Azithromycin |
| | Sánchez-Busó *et al*. 2019 (64) | 403 | PRJEB4024 | Azithromycin, Ceftriaxone, Cefixime, Ciprofloxacin, Tetracycline, Benzylpenicillin |
| | Grad *et al*. 2014, 2016 (18, 65) | 1,114 | PRJEB2090 PRJEB2999 PRJEB7904 | Azithromycin, Ceftriaxone, Cefixime, Ciprofloxacin, Tetracycline, Benzylpenicillin |
| | Jacobsson *et al*. 2016 (66) | 74 | PRJNA322768 | Azithromycin, Ceftriaxone, Cefixime, Ciprofloxacin |
| | Lee *et al*. 2018 (67) | 398 | PRJNA394216 | Azithromycin, Ceftriaxone, Cefixime, Ciprofloxacin, Tetracycline, Benzylpenicillin |
| | Unemo *et al*. 2016 (22) | 14 | PRJEB4024 | Azithromycin, Ceftriaxone, Cefixime, Ciprofloxacin, Tetracycline, Benzylpenicillin |
| Validation (N=1,607) | Town *et al*. 2020 (68, 117) | 1,288 | PRJEB19989 | Azithromycin, Ceftriaxone, Cefixime, Ciprofloxacin, Benzylpenicillin |
| | Yahara *et al*. 2018 (69) | 245 | PRJDB6496 PRJDB6504 | Azithromycin, Ceftriaxone, Cefixime, Ciprofloxacin |
| | Kwong *et al*. 2018 (70) | 75 | PRJEB17738 | Azithromycin, Ceftriaxone, Ciprofloxacin, Tetracycline, Benzylpenicillin, Spectinomycin |

Table S2. Point mutations and genes associated with antimicrobial resistance (AMR) detected by Pathogenwatch on the WHO 2016 reference genome panel. Note that screening for a *porA* mutant gene is included.

| Genome Name | Point mutations (SNPs and indels) | Genes |
|---|---|---|
| WHO_F | - | - |
| WHO_G | *folP*_R228S,*gyrA*_S91F,*mtrR*_promoter_a-57del,*parE*_G410V,*penA*_ins346D,*ponA1*_L421P, *rpsJ*_V57M | *tetM* |
| WHO_K | *folP*_R228S,*gyrA*_D95N,*gyrA*_S91F,*mtrR*_G45D,*mtrR*_promoter_a-57del,*parC*_S87R, *parC*_S88P,*penA*_G545S,*penA*_I312M,*penA*_V316T,*ponA1*_L421P,*porB1b*_A121D, *porB1b*_G120K,*rpsJ*_V57M | - |
| WHO_L | *gyrA*_D95N,*gyrA*_S91F,*mtrR*_G45D,*mtrR*_promoter_g-131a,*parC*_D86N,*parC*_S88P, *penA*_A501V,*penA*_G542S,*penA*_ins346D,*ponA1*_L421P,*porB1b*_A121D,*porB1b*_G120K, *rpsJ*_V57M | - |
| WHO_M | *folP*_R228S,*gyrA*_D95G,*gyrA*_S91F,*mtrR*_G45D,*mtrR*_promoter_a-57del,*penA*_ins346D, *ponA1*_L421P,*porB1b*_A121D,*porB1b*_G120K,*rpsJ*_V57M | *bla*TEM |
| WHO_N | *folP*_R228S,*gyrA*_D95G,*gyrA*_S91F,*mtrR*_A39T,*mtrR*_disrupted,*parC*_S87I,*parE*_G410V, *penA*_ins346D,*ponA1*_L421P,*rpsJ*_V57M | *bla*TEM,*tetM* |
| WHO_O | 16S_rDNA_c1184t,*folP*_R228S,*mtrR*_promoter_a-57del,*penA*_ins346D,*penA*_P551S, *ponA1*_L421P,*porB1b*_A121D,*porB1b*_G120K,*rpsJ*_V57M | *bla*TEM |
| WHO_P | *folP*_R228S,*mtrR*_disrupted,*penA*_ins346D,*porB1b*_A121D,*rpsJ*_V57M | *mtr*_mosaic_1, *mtrD*_mosaic_1 |
| WHO_U | 23S_rDNA_c2597t,*folP*_R228S,*penA*_ins346D,*ponA1*_L421P,*rpsJ*_V57M | *porA* |
| WHO_V | 23S_rDNA_a2045g,*folP*_R228S,*gyrA*_D95G,*gyrA*_S91F,*mtrR*_promoter_a-57del,*parC*_S87R, *penA*_G542S,*penA*_ins346D,*ponA1*_L421P,*porB1b*_A121D,*porB1b*_G120K,*rpsJ*_V57M | *bla*TEM |
| WHO_W | *folP*_R228S,*gyrA*_D95N,*gyrA*_S91F,*mtrR*_G45D,*mtrR*_promoter_a-57del,*parC*_S87R, *parC*_S88P,*penA*_G545S,*penA*_I312M,*penA*_V316T,*ponA1*_L421P,*porB1b*_A121D, *porB1b*_G120K,*rpsJ*_V57M | - |
| WHO_X | *folP*_R228S,*gyrA*_D95N,*gyrA*_S91F,*mtrR*_promoter_a-57del,*parC*_S87R,*parC*_S88P, *penA*_A311V,*penA*_G545S,*penA*_I312M,*penA*_T483S,*penA*_V316P,*ponA1*_L421P, *porB1b*_A121D,*porB1b*_G120K,*rpsJ*_V57M | - |
| WHO_Y | *folP*_R228S,*gyrA*_D95G,*gyrA*_S91F,*mtrR*_promoter_a-57del,*parC*_S87R,*penA*_A501P, *penA*_G545S,*penA*_I312M,*penA*_V316T,*ponA1*_L421P,*porB1b*_A121N,*porB1b*_G120K, *rpsJ*_V57M | - |
| WHO_Z | *folP*_R228S,*gyrA*_D95N,*gyrA*_S91F,*mtrR*_promoter_a-56c,*parC*_S87R,*parC*_S88P, *penA*_A311V,*penA*_G545S,*penA*_I312M,*penA*_T483S,*penA*_V316T,*ponA1*_L421P, *porB1b*_A121D,*porB1b*_G120K,*rpsJ*_V57M | - |

Table S3. Summary of the benchmark analysis of the list of genetic antimicrobial resistance (AMR) mechanisms. AZM = Azithromycin, CIP = Ciprofloxacin, CFM = Cefixime, CRO = Ceftriaxone, PEN = Benzylpenicillin, TET = Tetracycline, SPT = Spectinomycin, TP = True Positives, FP = False Positives, TN = True Negatives, FN = False Negatives, NPV = Negative Predictive Value, CI = Confidence Intervals.

| Antibiotic | Dataset | TOTAL | TP | FP | TN | FN | Sensitivity | Specificity | NPV | PPV |
|---|---|---|---|---|---|---|---|---|---|---|
| AZM | Test (CI) | 3679 | 468 | 17 | 3008 | 186 | 71.56 (67.93-74.99) | 99.44 (99.10-99.67) | 94.18 (93.31-94.96) | 96.49 (94.45-97.95) |
| | Validation (CI) | 1573 | 8 | 7 | 1556 | 2 | 80.00 (44.39-97.48) | 99.55 (99.08-99.82) | 99.87 (99.54-99.98) | 53.33 (26.59-78.73) |
| CFM | Test (CI) | 3601 | 371 | 323 | 2892 | 15 | 96.11 (93.67-97.81) | 89.95 (88.86-90.97) | 99.48 (99.15-99.71) | 53.46 (49.67-57.22) |
| | Validation (CI) | 1498 | 68 | 137 | 1289 | 4 | 94.44 (86.38-98.47) | 90.39 (88.74-91.87) | 99.69 (99.21-99.92) | 33.17 (26.77-40.07) |
| CIP | Test (CI) | 3281 | 1548 | 15 | 1671 | 47 | 97.05 (96.10-97.83) | 99.11 (98.54-99.50) | 97.26 (96.38-97.98) | 99.04 (98.42-99.46) |
| | Validation (CI) | 1290 | 549 | 4 | 693 | 44 | 92.58 (90.17-94.56) | 99.43 (98.54-99.84) | 94.03 (92.07-95.63) | 99.28 (98.16-99.80) |
| CRO | Test (CI) | 3635 | 9 | 5 | 3603 | 18 | 33.33 (16.52-53.96) | 99.86 (99.68-99.95) | 99.50 (99.22-99.71) | 64.29 (35.14-87.24) |
| | Validation (CI) | 1571 | 3 | 0 | 1565 | 3 | 50.00 (11.81-88.19) | 100.00 (99.76-100) | 99.81 (99.44-99.96) | 100.00 (29.24-100) |
| PEN | Test (CI) | 1654 | 1424 | 46 | 157 | 27 | 98.14 (97.30-98.77) | 77.34 (70.96-82.91) | 85.33 (79.37-90.10) | 96.87 (95.85-97.70) |
| | Validation (CI) | 1330 | 1228 | 86 | 7 | 9 | 99.27 (98.62-99.67) | 7.53 (3.08-14.90) | 43.75 (19.75-70.12) | 93.46 (91.98-94.73) |
| TET | Test (CI) | 1661 | 1096 | 215 | 341 | 9 | 99.19 (98.46-99.63) | 61.33 (57.14-65.40) | 97.43 (95.17-98.82) | 83.60 (81.48-85.57) |
| | Validation (CI) | 75 | 74 | 0 | 0 | 1 | 98.67 (92.79-99.97) | - | 0.00 (0.00-97.50) | 100.00 (95.14-100) |
| SPT | Validation (CI) | 75 | 0 | 0 | 75 | 0 | - | 100.00 (95.20-100) | 100.00 (95.20-100) | - |

Table S4. List of genetic mechanisms detected on the test benchmark (N=3,987) for each of the six main antibiotics. The total number of isolates carrying each mechanism and the benchmark results are shown. AZM = Azithromycin, CIP = Ciprofloxacin, CFM = Cefixime, CRO = Ceftriaxone, PEN = Benzylpenicillin, TET = Tetracycline, SPT = Spectinomycin, TP = True Positives, FP = False Positives, PPV = Positive Predictive Values.

| Agent | Mechanism | Total | TP | FP | PPV |
|---|---|---|---|---|---|
| AZM | 23S_rDNA_a2045g | 108 | 106 | 2 | 98.15 |
| AZM | 23S_rDNA_c2597t | 340 | 331 | 9 | 97.35 |
| AZM | 23S_rDNA_c2597t__*mtrR*_promoter_a-57del__*mtrR*_G45D | 6 | 6 | 0 | 100 |
| AZM | *ermB* | 1 | 1 | 0 | 100 |
| AZM | *ermC* | 2 | 2 | 0 | 100 |
| AZM | *mtr*_mosaic_1 | 6 | 4 | 2 | 66.67 |
| AZM | *mtr*_mosaic_2 | 24 | 21 | 3 | 87.5 |
| AZM | *mtr*_mosaic_3 | 1 | 1 | 0 | 100 |
| AZM | *mtrD*_mosaic_1 | 8 | 5 | 3 | 62.5 |
| AZM | *mtrD*_mosaic_2 | 21 | 19 | 2 | 90.48 |
| AZM | *mtrD*_mosaic_3 | 1 | 1 | 0 | 100 |
| AZM | *rplD*_G70D__23S_rDNA_c2597t | 5 | 5 | 0 | 100 |
| AZM | *rplV*_------83KGPSLK | 1 | 1 | 0 | 100 |
| AZM | *rplV*_----90ARAK | 1 | 1 | 0 | 100 |
| CFM | *penA*_A311V_G545S_I312M_T483S_V316P | 1 | 1 | 0 | 100 |
| CFM | *penA*_A311V_G545S_I312M_T483S_V316T | 1 | 1 | 0 | 100 |
| CFM | *penA*_A501P | 4 | 4 | 0 | 100 |
| CFM | *penA*_G545S_I312M_V316T | 690 | 367 | 323 | 53.19 |
| CFM | *penA*_G545S_I312M_V316T__*mtrR*_G45D | 50 | 39 | 11 | 78 |
| CFM | *penA*_T483S | 2 | 2 | 0 | 100 |
| CFM | *penA*_V316P | 1 | 1 | 0 | 100 |
| CFM | *rpoB*_R201H | 1 | 1 | 0 | 100 |
| CFM | *rpoB*_R201H__*mtrR*_G45D | 1 | 1 | 0 | 100 |
| CFM | *rpoD*_A95-_D92-_D93-_D94- | 1 | 1 | 0 | 100 |
| CFM | *rpoD*_A95-_D92-_D93-_D94-__*mtrR*_G45D | 1 | 1 | 0 | 100 |
| CFM | *rpoD*_E98K | 1 | 1 | 0 | 100 |
| CFM | *rpoD*_E98K__*mtrR*_G45D | 1 | 1 | 0 | 100 |
| CIP | *gyrA*_D95A | 227 | 225 | 2 | 99.12 |
| CIP | *gyrA*_D95G | 1199 | 1192 | 7 | 99.42 |
| CIP | *gyrA*_D95N | 57 | 57 | 0 | 100 |
| CIP | *gyrA*_S91F | 1511 | 1503 | 8 | 99.47 |
| CIP | *parC*_D86N | 90 | 90 | 0 | 100 |
| CIP | *parC*_E91K | 22 | 22 | 0 | 100 |
| CIP | *parC*_S87I | 14 | 14 | 0 | 100 |
| CIP | *parC*_S87N | 98 | 98 | 0 | 100 |
| CIP | *parC*_S87R | 1050 | 1037 | 13 | 98.76 |
| CIP | *parC*_S88P | 25 | 25 | 0 | 100 |
| CIP | *parE*_G410V | 5 | 5 | 0 | 100 |
| CRO | *penA*_A311V_G545S_I312M_T483S_V316P | 1 | 1 | 0 | 100 |
| CRO | *penA*_A311V_G545S_I312M_T483S_V316T | 1 | 1 | 0 | 100 |
| CRO | *penA*_A501P | 4 | 4 | 0 | 100 |
| CRO | *penA*_A501V_G542S | 5 | 2 | 3 | 40 |
| CRO | *penA*_T483S | 2 | 2 | 0 | 100 |
| CRO | *penA*_V316P | 1 | 1 | 0 | 100 |
| CRO | *rpoB*_R201H | 1 | 1 | 0 | 100 |
| CRO | *rpoD*_A95-_D92-_D93-_D94- | 1 | 0 | 1 | 0 |
| CRO | *rpoD*_E98K | 1 | 0 | 1 | 0 |
| PEN | *blaTEM* | 133 | 129 | 4 | 96.99 |
| PEN | *mtrR*_A39T__*porB1b*_A121D_G120K | 31 | 31 | 0 | 100 |
| PEN | *mtrR*_disrupted | 67 | 65 | 2 | 97.01 |
| PEN | *mtrR*_G45D | 231 | 218 | 13 | 94.37 |
| PEN | *mtrR*_promoter_a-56c | 1 | 1 | 0 | 100 |
| PEN | *mtrR*_promoter_a-57del | 687 | 665 | 22 | 96.8 |
| PEN | *mtrR*_promoter_a-57del__*penA*_G542S__*porB1b*_A121D_G120K | 42 | 41 | 1 | 97.62 |
| PEN | *mtrR*_promoter_a-57del__*porB1b*_A121N_G120K | 286 | 276 | 10 | 96.5 |
| PEN | *mtrR*_promoter_g-131a | 9 | 9 | 0 | 100 |
| PEN | *penA*_A501P | 4 | 4 | 0 | 100 |
| PEN | *penA*_A501T | 55 | 55 | 0 | 100 |
| PEN | *penA*_A501V | 30 | 30 | 0 | 100 |
| PEN | *penA*_G542S | 120 | 118 | 2 | 98.33 |
| PEN | *penA*_G545S | 297 | 289 | 8 | 97.31 |
| PEN | *penA*_G545S_I312M_V316T__*porB1b*_A121D_G120K | 17 | 17 | 0 | 100 |
| PEN | *penA*_I312M | 316 | 306 | 10 | 96.84 |
| PEN | *penA*_ins346D | 1124 | 1091 | 33 | 97.06 |
| PEN | *penA*_P551S | 92 | 87 | 5 | 94.57 |
| PEN | *penA*_T483S | 2 | 2 | 0 | 100 |
| PEN | *penA*_V316P | 1 | 1 | 0 | 100 |
| PEN | *penA*_V316T | 315 | 305 | 10 | 96.83 |

| | | | | | |
|---|---|---|---|---|---|
| PEN | *ponA1*_L421P | 786 | 764 | 22 | 97.2 |
| PEN | *porB1b*_A121D | 323 | 314 | 9 | 97.21 |
| PEN | *porB1b*_A121D__*ponA1*_L421P__*mtrR*_G45D__*mtrR*_promoter_a-57del | 34 | 33 | 1 | 97.06 |
| PEN | *porB1b*_A121N | 305 | 293 | 12 | 96.07 |
| PEN | *porB1b*_A121N_G120K__*ponA1*_L421P__*mtrR*_promoter_a-57del | 269 | 259 | 10 | 96.28 |
| PEN | *porB1b*_G120K | 618 | 597 | 21 | 96.6 |
| TET | *mtrR*_disrupted | 67 | 62 | 5 | 92.54 |
| TET | *mtrR*_promoter_a-56c | 1 | 1 | 0 | 100 |
| TET | *mtrR*_promoter_a-57del | 691 | 651 | 40 | 94.21 |
| TET | *mtrR*_promoter_a-57del__*rpsJ*_V57M | 681 | 646 | 35 | 94.86 |
| TET | *mtrR*_promoter_a-57del__*rpsJ*_V57M__*mtrR*_G45D | 83 | 78 | 5 | 93.98 |
| TET | *mtrR*_promoter_g-131a | 9 | 7 | 2 | 77.78 |
| TET | *rpsJ*_V57M | 1292 | 1083 | 209 | 83.82 |
| TET | *rpsJ*_V57M__*mtrR*_A39T__*mtrR*_promoter_a-57del | 6 | 5 | 1 | 83.33 |
| TET | *rpsJ*_V57M__*mtrR*_A39T_disrupted | 51 | 51 | 0 | 100 |
| TET | *rpsJ*_V57M__*mtrR*_A39T_G45D | 3 | 3 | 0 | 100 |
| TET | *rpsJ*_V57M__*mtrR*_promoter_a-56c | 1 | 1 | 0 | 100 |
| TET | *tetM* | 245 | 243 | 2 | 99.18 |

Table S5. List of genetic mechanisms detected on the validation benchmark (N=1,607) for each of the six main antibiotics. The total number of isolates carrying each mechanism and the benchmark results are shown. AZM = Azithromycin, CIP = Ciprofloxacin, CFM = Cefixime, CRO = Ceftriaxone, PEN = Benzylpenicillin, TET = Tetracycline, SPT = Spectinomycin, TP = True Positives, FP = False Positives, PPV = Positive Predictive Values.

| Agent | Mechanism | Total | TP | FP | PPV |
|---|---|---|---|---|---|
| AZM | 23S_rDNA_a2045g | 3 | 3 | 0 | 100 |
| AZM | 23S_rDNA_c2597t | 3 | 3 | 0 | 100 |
| AZM | *mtr*_mosaic_1 | 2 | 1 | 1 | 50 |
| AZM | *mtr*_mosaic_2 | 8 | 2 | 6 | 25 |
| AZM | *mtrD*_mosaic_1 | 1 | 1 | 0 | 100 |
| AZM | *mtrD*_mosaic_2 | 6 | 2 | 4 | 33.33 |
| CFM | *penA*_A311V_G545S_I312M_T483S_V316T | 3 | 3 | 0 | 100 |
| CFM | *penA*_G545S_I312M_V316T | 205 | 68 | 137 | 33.17 |
| CFM | *penA*_G545S_I312M_V316T__*mtrR*_G45D | 32 | 21 | 11 | 65.63 |
| CFM | *penA*_T483S | 3 | 3 | 0 | 100 |
| CIP | *gyrA*_D95A | 188 | 185 | 3 | 98.4 |
| CIP | *gyrA*_D95G | 309 | 308 | 1 | 99.68 |
| CIP | *gyrA*_D95N | 55 | 55 | 0 | 100 |
| CIP | *gyrA*_S91F | 549 | 545 | 4 | 99.27 |
| CIP | *parC*_D86N | 90 | 90 | 0 | 100 |
| CIP | *parC*_E91K | 4 | 4 | 0 | 100 |
| CIP | *parC*_S87I | 6 | 6 | 0 | 100 |
| CIP | *parC*_S87N | 31 | 30 | 1 | 96.77 |
| CIP | *parC*_S87R | 321 | 320 | 1 | 99.69 |
| CIP | *parC*_S88P | 44 | 44 | 0 | 100 |
| CRO | *penA*_A311V_G545S_I312M_T483S_V316T | 3 | 3 | 0 | 100 |
| CRO | *penA*_T483S | 3 | 3 | 0 | 100 |
| PEN | *blaTEM* | 561 | 525 | 36 | 93.58 |
| PEN | *mtrR*_A39T__*porB1b*_A121D_G120K | 11 | 11 | 0 | 100 |
| PEN | *mtrR*_disrupted | 94 | 91 | 3 | 96.81 |
| PEN | *mtrR*_G45D | 144 | 141 | 3 | 97.92 |
| PEN | *mtrR*_promoter_a-56c | 2 | 2 | 0 | 100 |
| PEN | *mtrR*_promoter_a-57del | 395 | 388 | 7 | 98.23 |
| PEN | *mtrR*_promoter_a-57del__*penA*_G542S__*porB1b*_A121D_G120K | 2 | 2 | 0 | 100 |
| PEN | *mtrR*_promoter_a-57del__*porB1b*_A121N_G120K | 60 | 60 | 0 | 100 |
| PEN | *mtrR*_promoter_g-131a | 3 | 3 | 0 | 100 |
| PEN | *penA*_A501T | 98 | 98 | 0 | 100 |
| PEN | *penA*_A501V | 30 | 30 | 0 | 100 |
| PEN | *penA*_G542S | 31 | 31 | 0 | 100 |
| PEN | *penA*_G545S | 93 | 93 | 0 | 100 |
| PEN | *penA*_G545S_I312M_V316T__*porB1b*_A121D_G120K | 2 | 2 | 0 | 100 |
| PEN | *penA*_I312M | 94 | 94 | 0 | 100 |
| PEN | *penA*_ins346D | 1208 | 1125 | 83 | 93.13 |
| PEN | *penA*_P551S | 42 | 42 | 0 | 100 |
| PEN | *penA*_V316T | 94 | 94 | 0 | 100 |
| PEN | *ponA1*_L421P | 464 | 455 | 9 | 98.06 |
| PEN | *porB1b*_A121D | 192 | 189 | 3 | 98.44 |
| PEN | *porB1b*_A121D__*ponA1*_L421P__*mtrR*_G45D__*mtrR*_promoter_a-57del | 11 | 10 | 1 | 90.91 |
| PEN | *porB1b*_A121N | 83 | 83 | 0 | 100 |
| PEN | *porB1b*_A121N_G120K__*ponA1*_L421P__*mtrR*_promoter_a-57del | 60 | 60 | 0 | 100 |
| PEN | *porB1b*_G120K | 283 | 281 | 2 | 99.29 |
| TET | *mtrR*_disrupted | 2 | 2 | 0 | 100 |
| TET | *mtrR*_promoter_a-57del | 37 | 37 | 0 | 100 |
| TET | *mtrR*_promoter_a-57del__*rpsJ*_V57M | 37 | 37 | 0 | 100 |
| TET | *mtrR*_promoter_a-57del__*rpsJ*_V57M__*mtrR*_G45D | 3 | 3 | 0 | 100 |
| TET | *mtrR*_promoter_g-131a | 3 | 3 | 0 | 100 |
| TET | *rpsJ*_V57M | 74 | 74 | 0 | 100 |
| TET | *rpsJ*_V57M__*mtrR*_A39T_disrupted | 2 | 2 | 0 | 100 |
| TET | *tetM* | 5 | 5 | 0 | 100 |

Table S6. Published studies on *N. gonorrhoeae* genomics for which new public collections have been created in Pathogenwatch sorted by included number of isolates. Dates correspond to those in the final collections.

| PubMed ID (reference) | ENA project accession(s) | Collection label (reference) | Number of isolates in publication | Number of isolates in collection | Geographical location and timescale |
|---|---|---|---|---|---|
| 31488838 | PRJNA520805 | Williamson et al, 2019 (118) | 2,186 | 2,179 | Australia<br>2017 |
| 27427203 | PRJNA315363 | De Silva et al, 2016 (119) | 1,872 | 1,783 | Brighton, United Kingdom (UK)<br>2004-2015 |
| 31978353, 32091356 | PRJEB19989 | Town et al, 2020* (68, 117) | 1,277 | 1,288 | England, UK<br>2013-2016 |
| 27638945 | PRJEB2090<br>PRJEB2999<br>PRJEB7904 | Grad et al, 2016* (18) | 1,102 | 1,035 | United States (US)<br>2000-2013 |
| 32829411 | PRJEB10016 | Mortimer et al, 2020 (120) | 897 | 891 | New York City, US<br>2011-2015 |
| 32213251 | PRJEB32435 | Alfsnes et al, 2020 (121) | 958 | 816 | Norway<br>2016-2017 |
| 30788502 | PRJNA317462<br>PRJNA329501 | Thomas et al, 2019 (122) | 649 | 644 | United States<br>2014-2016 |
| 31358980 | PRJEB4024 | Sánchez-Busó et al, 2019 (64) | 419 | 395 | Worldwide<br>1979-2013 |
| 29182725 | PRJNA394216 | Lee et al, 2018 (67) | 398 | 376 | New Zealand<br>2014-2015 |
| 32071056 | PRJNA317462<br>PRJNA329501 | Schmerer et al, 2020 (123) | 334 | 324 | United States<br>2016 |
| 30063202 | PRJDB6496<br>PRJDB6504 | Yahara et al, 2018 (69) | 271 | 245 | Kyoto and Osaka, Japan<br>2011-2015 |
| 32068837 | PRJEB34425 | Lan et al, 2020 (124) | 229 | 227 | Vietnam<br>2011-2016 |
| 24462211 | PRJEB2090<br>PRJEB2999<br>PRJEB7904 | Grad et al, 2014* (65) | 236 | 216 | United States<br>2009-2010 |
| 26935729 | PRJNA298332 | Demczuk et al, 2016 (61) | 246 | 200 | Canada<br>1997-2014 |
| 27353752 | PRJEB2124 | Didelot et al, 2016 (125) | 237 | 194 | Sheffield and London, UK<br>1995-2004 |
| 32013864 | PRJEB4024 | Golparian et al, 2020 (59) | 231 | 192 | Denmark<br>1928-2013 |
| 25378573 | PRJNA266539 | Demczuk et al, 2015 (60) | 180 | 168 | Canada<br>1989-2013 |
| 33200978 | PRJEB32435 | Osnes et al, 2020 (126) | 148 | 133 | Norway<br>2015-2018 |
| 29701830 | PRJEB10104 | Cehovin et al, 2018* (127) | 103 | 112 | Coastal Kenya<br>2010-2015 |
| 29523496 | PRJEB23008 | Fifer et al, 2018 (63) | 101 | 100 | England, UK<br>2004-2017 |
| 29367612 | PRJNA392203 | Buckley et al, 2018 (128) | 94 | 92 | Australia<br>2012-2014 |
| 29247013 | PRJEB17738 | Kwong et al, 2018 (70) | 94 | 75 | Australia<br>2006-2014 |

| | | | | | |
|---|---|---|---|---|---|
| 28348871 | PRJEB14168 | Kwong et al, 2016 (53); Martin et al, 2004 (49) | 50 | 48 | Australia and New Zealand 2004-2015 |
| 29882175 | PRJNA473385 | Ryan et al, 2018 (129) | 43 | 42 | Ireland 2012-2016 |
| 28510723 | PRJNA348107 | Wind et al, 2017 (130) | 31 | 23 | Amsterdam, Netherlands 2002-2012 |
| 25780762 | ** | Ezewudo et al, 2015 (131) | 61 | 18 | Worldwide 1982-2008 |
| 26601852 | PRJEB14933 | Chisholm et al, 2015 (58) | 15 | 14 | Leeds, UK 2015 |

* In these collections, the number of accession numbers on the corresponding ENA project was higher than the number in the final publication. These genomes were included in the corresponding collections.
** Multiple ENA project accessions are linked to Ezewudo *et al*, 2015: PRJNA209340, PRJNA209307, PRJNA209319, PRJNA209352, PRJNA209376, PRJNA209466, PRJNA209373, PRJNA209465, PRJNA209320, PRJNA209333, PRJNA209345, PRJNA209351, PRJNA209342, PRJNA209343, PRJNA209347, PRJNA209470, PRJNA244850, PRJNA209316.

Table S7. Number of public *N. gonorrhoeae* genomes in Pathogenwatch clustered by country.

| Country | Number of isolates | Individual proportion (%) | Cumulative sum | Cumulative proportion (%) |
|---|---|---|---|---|
| United Kingdom | 3476 | 27.77 | 3476 | 27.77 |
| United States | 2774 | 22.17 | 6250 | 49.94 |
| Australia | 2388 | 19.08 | 8638 | 69.02 |
| Norway | 990 | 7.91 | 9628 | 76.93 |
| New Zealand | 396 | 3.16 | 10024 | 80.10 |
| Canada | 382 | 3.05 | 10406 | 83.15 |
| Japan | 268 | 2.14 | 10674 | 85.29 |
| Denmark | 255 | 2.04 | 10929 | 87.33 |
| Vietnam | 233 | 1.86 | 11162 | 89.19 |
| Spain | 129 | 1.03 | 11291 | 90.22 |
| Kenya | 112 | 0.89 | 11403 | 91.11 |
| Portugal | 108 | 0.86 | 11511 | 91.98 |
| Netherlands | 92 | 0.74 | 11603 | 92.71 |
| Slovenia | 77 | 0.62 | 11680 | 93.33 |
| France | 62 | 0.50 | 11742 | 93.82 |
| Belgium | 55 | 0.44 | 11797 | 94.26 |
| Austria | 54 | 0.43 | 11851 | 94.69 |
| Greece | 54 | 0.43 | 11905 | 95.13 |
| Germany | 53 | 0.42 | 11958 | 95.55 |
| Sweden | 51 | 0.41 | 12009 | 95.96 |
| Hungary | 48 | 0.38 | 12057 | 96.34 |
| Ireland | 42 | 0.34 | 12099 | 96.68 |
| Slovakia | 39 | 0.31 | 12138 | 96.99 |
| Latvia | 38 | 0.30 | 12176 | 97.29 |
| Poland | 34 | 0.27 | 12210 | 97.56 |
| Scotland | 30 | 0.24 | 12240 | 97.80 |
| Italy | 28 | 0.22 | 12268 | 98.03 |
| Belarus | 24 | 0.19 | 12292 | 98.22 |
| India | 24 | 0.19 | 12316 | 98.41 |
| Guinea-Bissau | 22 | 0.18 | 12338 | 98.59 |
| Thailand | 22 | 0.18 | 12360 | 98.76 |
| Malta | 20 | 0.16 | 12380 | 98.92 |
| Estonia | 17 | 0.14 | 12397 | 99.06 |
| Pakistan | 14 | 0.11 | 12411 | 99.17 |
| Philippines | 14 | 0.11 | 12425 | 99.28 |
| Russia | 13 | 0.10 | 12438 | 99.38 |
| Cyprus | 8 | 0.06 | 12446 | 99.45 |
| Bhutan | 7 | 0.06 | 12453 | 99.50 |
| China | 6 | 0.05 | 12459 | 99.55 |
| Gambia | 5 | 0.04 | 12464 | 99.59 |
| Iceland | 5 | 0.04 | 12469 | 99.63 |
| Indonesia | 5 | 0.04 | 12474 | 99.67 |
| Cuba | 4 | 0.03 | 12478 | 99.70 |
| Turkey | 4 | 0.03 | 12482 | 99.74 |
| Morocco | 3 | 0.02 | 12485 | 99.76 |
| South Africa | 3 | 0.02 | 12488 | 99.78 |
| Brasil | 2 | 0.02 | 12490 | 99.80 |
| Cabo Verde | 2 | 0.02 | 12492 | 99.82 |
| Chile | 2 | 0.02 | 12494 | 99.83 |
| Ivory Coast | 2 | 0.02 | 12496 | 99.85 |
| Suriname | 2 | 0.02 | 12498 | 99.86 |
| Tanzania | 2 | 0.02 | 12500 | 99.88 |
| Angola | 1 | 0.01 | 12501 | 99.89 |
| Argentina | 1 | 0.01 | 12502 | 99.90 |
| Armenia | 1 | 0.01 | 12503 | 99.90 |
| Bulgaria | 1 | 0.01 | 12504 | 99.91 |
| Caribbean | 1 | 0.01 | 12505 | 99.92 |

| | | | | |
|---|---|---|---|---|
| Ecuador | 1 | 0.01 | 12506 | 99.93 |
| Finland | 1 | 0.01 | 12507 | 99.94 |
| Guinea | 1 | 0.01 | 12508 | 99.94 |
| Hong Kong | 1 | 0.01 | 12509 | 99.95 |
| Jamaica | 1 | 0.01 | 12510 | 99.96 |
| Lithuania | 1 | 0.01 | 12511 | 99.97 |
| Malaysia | 1 | 0.01 | 12512 | 99.98 |
| Romania | 1 | 0.01 | 12513 | 99.98 |
| Saudi Arabia | 1 | 0.01 | 12514 | 99.99 |
| Uganda | 1 | 0.01 | 12515 | 100.00 |
