## Additional file 3 - Supplementary Figures for "A community-driven resource for genomic epidemiology and antimicrobial resistance prediction of *Neisseria gonorrhoeae* at Pathogenwatch"

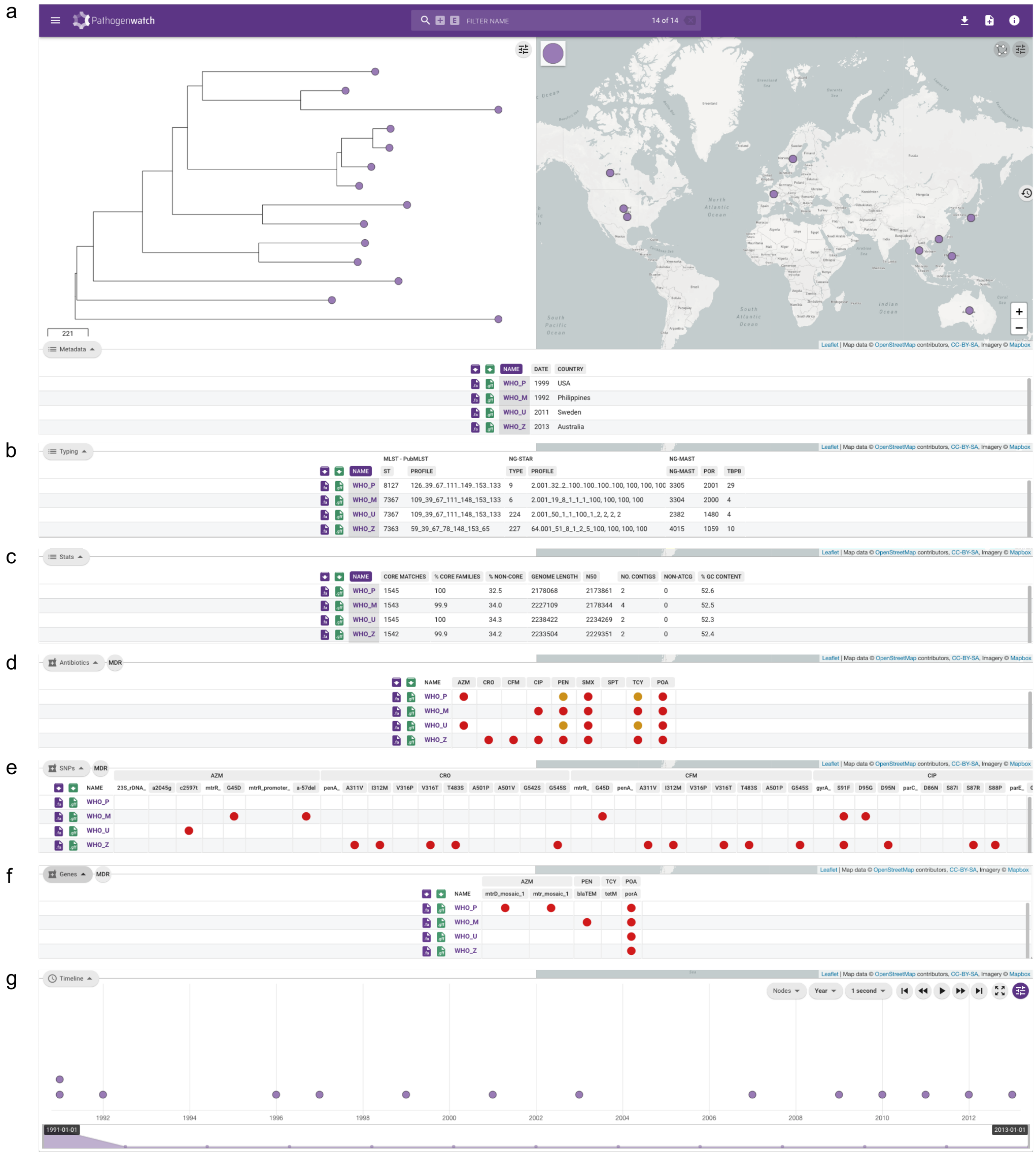


Figure S1. (a) Pathogenwatch layout showing the phylogenetic tree on the left, the map with the locations of isolation of each isolate and the metadata table. The latter can be switched among different options, such as the typing results (b), assembly statistics (c), genotypic prediction for antimicrobial resistance (d, orange for susceptibility but increased exposure and red for complete resistance), single nucleotide polymorphisms (SNPs) involved in antimicrobial resistance (AMR) (e), genes involved in AMR (f) and a timeline (g). This example shows the 14 WHO 2016 *N. gonorrhoeae* reference genomes (22).

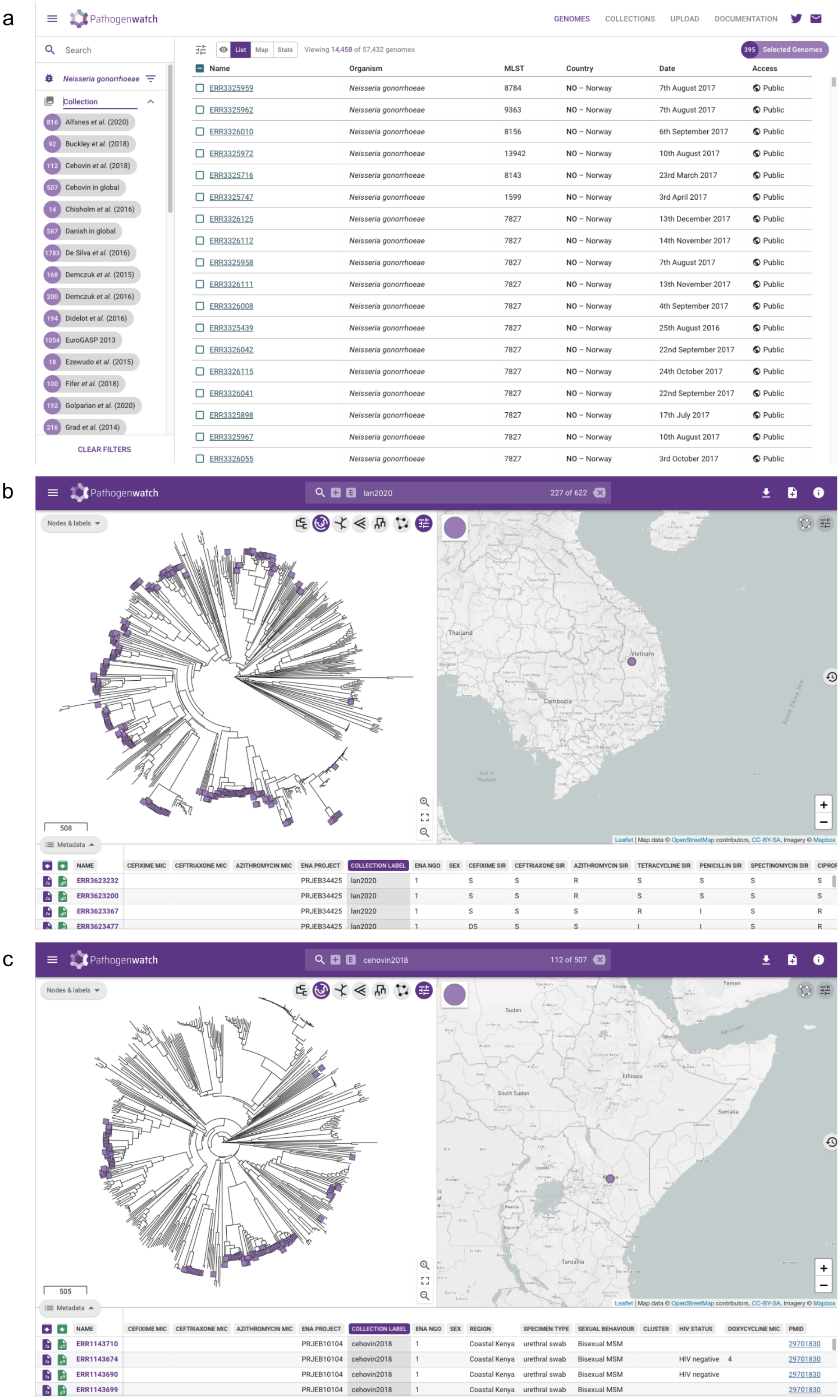


Figure S2. (a) In the genomes page, the users can combine genomes and collections in a ‘shopping cart’ mode. Pathogenwatch will join the metadata, typing and antimicrobial resistance (AMR) results and will create a combined phylogenetic tree. (b) Example of a collection of *N. gonorrhoeae* isolates from Vietnam (2011-2016) (124) put in context on a global collection (1979-2013) (64). (c) Example of a collection of *N. gonorrhoeae* isolates from Coastal Kenya (2020-2015) (127) put in context on the same global collection.

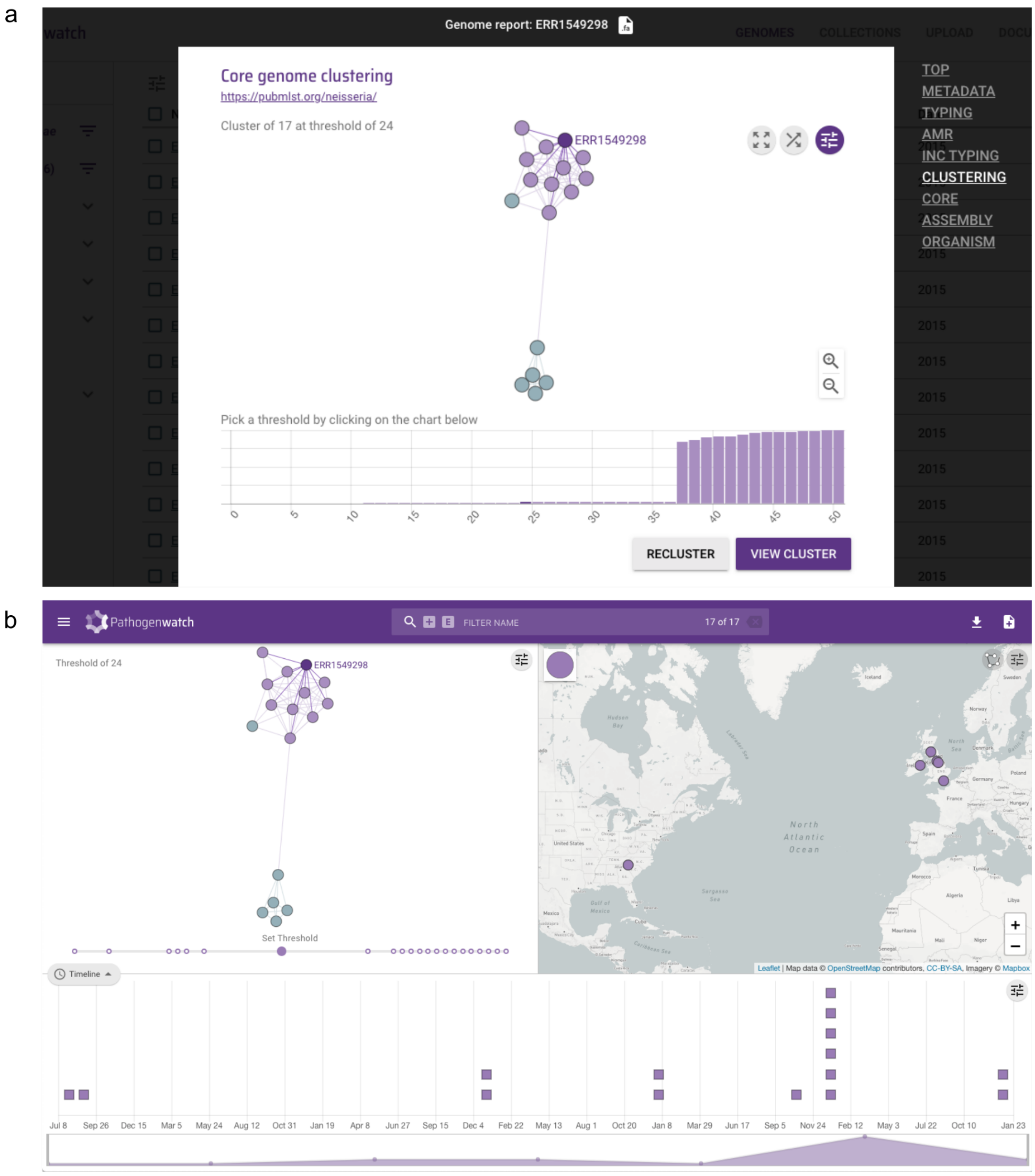


Figure S3. (a) From each individual genome report, users can access a cluster view that shows which other genomes in Pathogenwatch are close to it based on a threshold determined by the number of differing loci on the *N. gonorrhoeae* core genome MLST (cgMLST) v1.0 scheme (light purple dots). Light green dots correspond to genomes that are recursively linked to the light purple under the same chosen threshold. From the 'View Cluster' button on the report, the user can access a collection containing the genomes in the selected cluster (b).

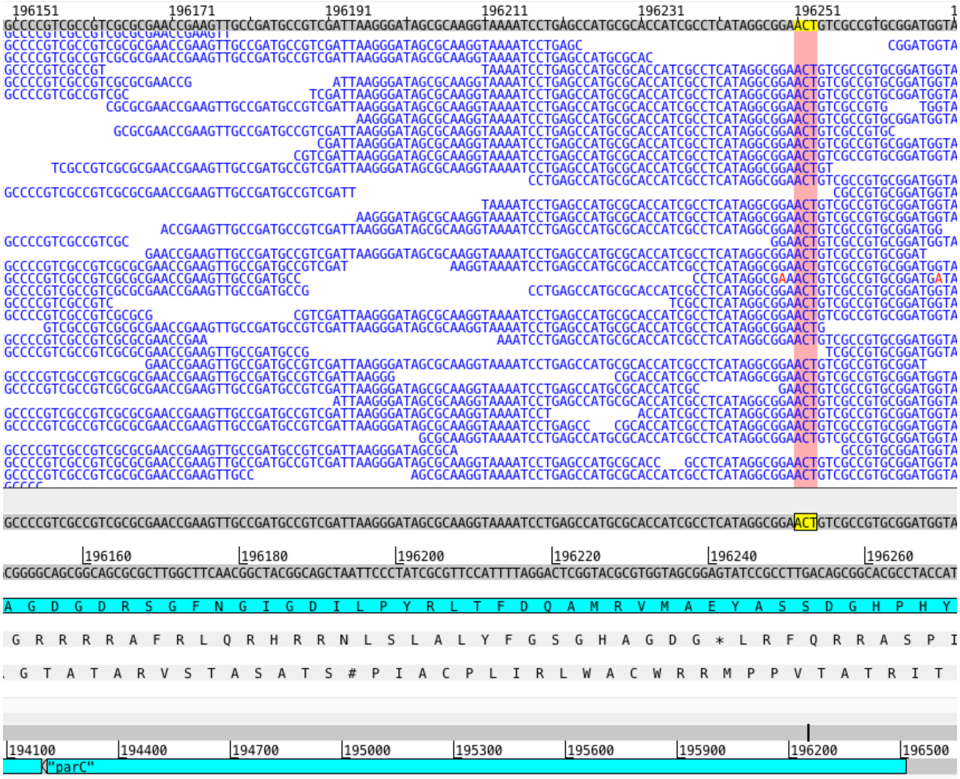

Figure S4. Artemis (73) visualization of Illumina short reads for the WHO U reference strain (European Nucleotide Archive (ENA) run accession ERR449479) mapped to the reference genome assembly (ENA genome accession LT592159.1). A snapshot of the *parC* gene is shown, with codon 87 (5'-TCA-3') highlighted in pink, corresponding to amino acid Serine (S).

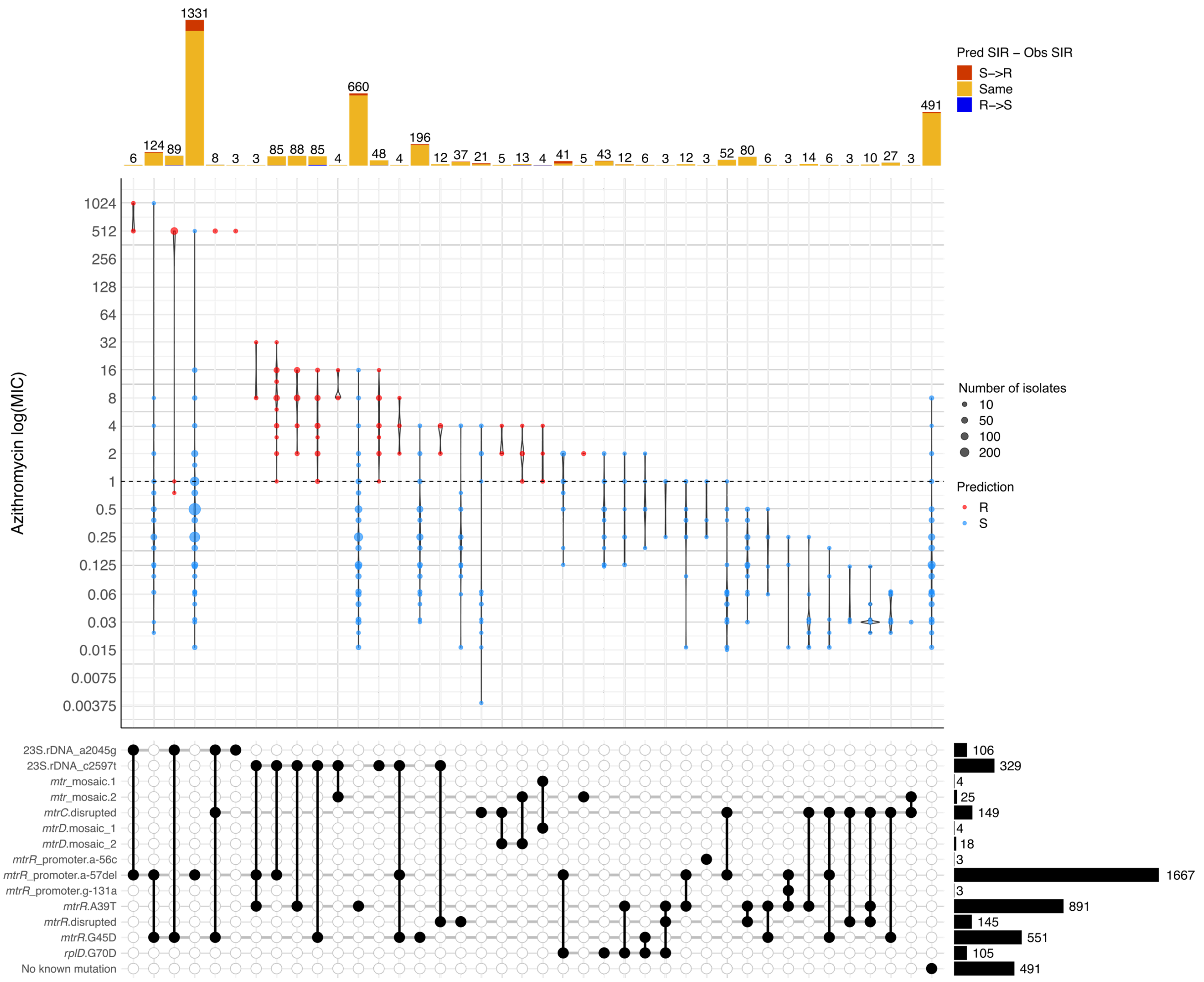


Figure S5. Distribution of minimum inhibitory concentration (MIC) (mg/L) values for azithromycin in a collection of 3,987 *N. gonorrhoeae* isolates with different combinations of antimicrobial resistance (AMR) genetic mechanisms. Combinations observed in at least 3 isolates are shown. Dashed horizontal lines on the violin plots mark the EUCAST epidemiological cut-off (ECOFF). Point colours inside violins represent the genotypic AMR prediction by Pathogenwatch on each combination of mechanisms. Barplots on the top show the abundance of isolates with each combination of mechanisms. Bar colours represent the differences between the predicted (Pred SIR) and the observed SIR (Obs SIR) (i.e. red for a predicted susceptible mechanism when the observed phenotype is resistant). Black bars on the bottom right represent the number of isolates with each particular mechanism individually or in combination.

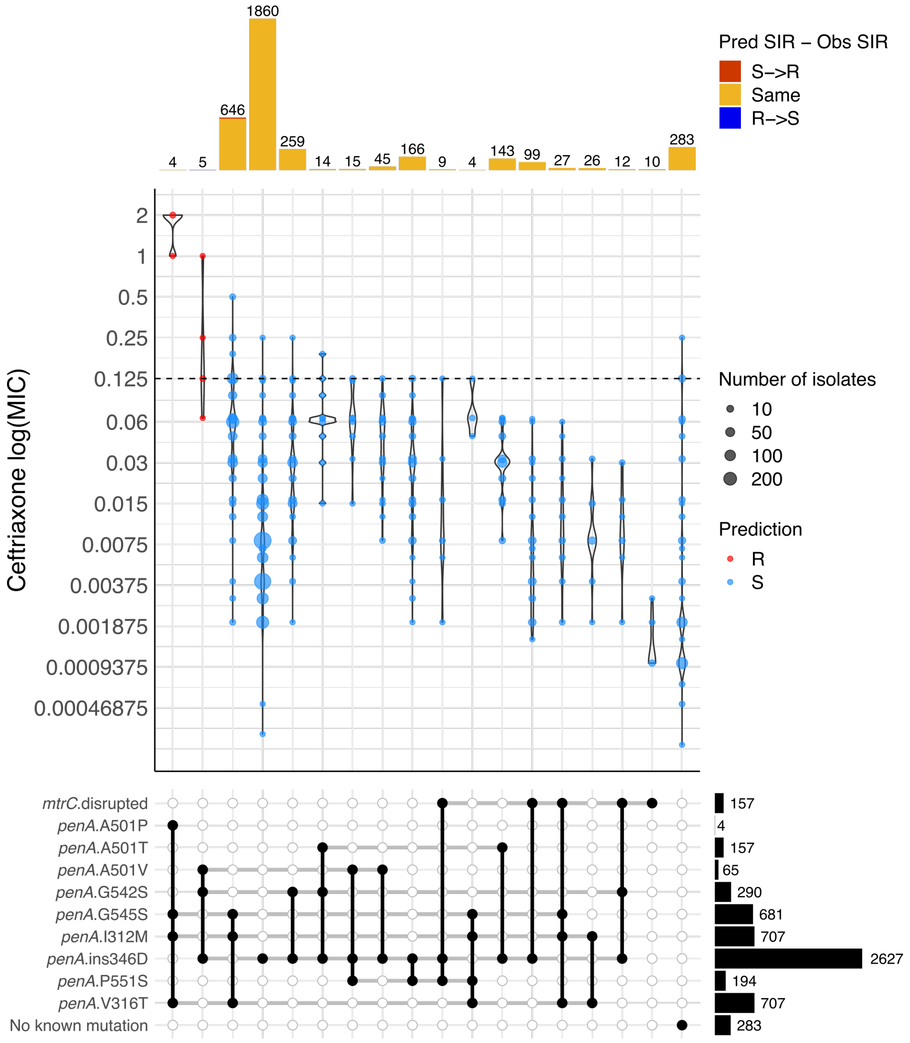


Figure S6. Distribution of minimum inhibitory concentration (MIC) (mg/L) values for ceftriaxone in a collection of 3,987 *N. gonorrhoeae* isolates with different combinations of antimicrobial resistance (AMR) genetic mechanisms. Combinations observed in at least 3 isolates are shown. Dashed horizontal lines on the violin plots mark the EUCAST resistance breakpoint. Point colours inside violins represent the genotypic AMR prediction by Pathogenwatch on each combination of mechanisms. Barplots on the top show the abundance of isolates with each combination of mechanisms. Bar colours represent the differences between the predicted (Pred SIR) and the observed SIR (Obs SIR) (i.e. red for a predicted susceptible mechanism when the observed phenotype is resistant). Black bars on the bottom right represent the number of isolates with each particular mechanism individually or in combination.

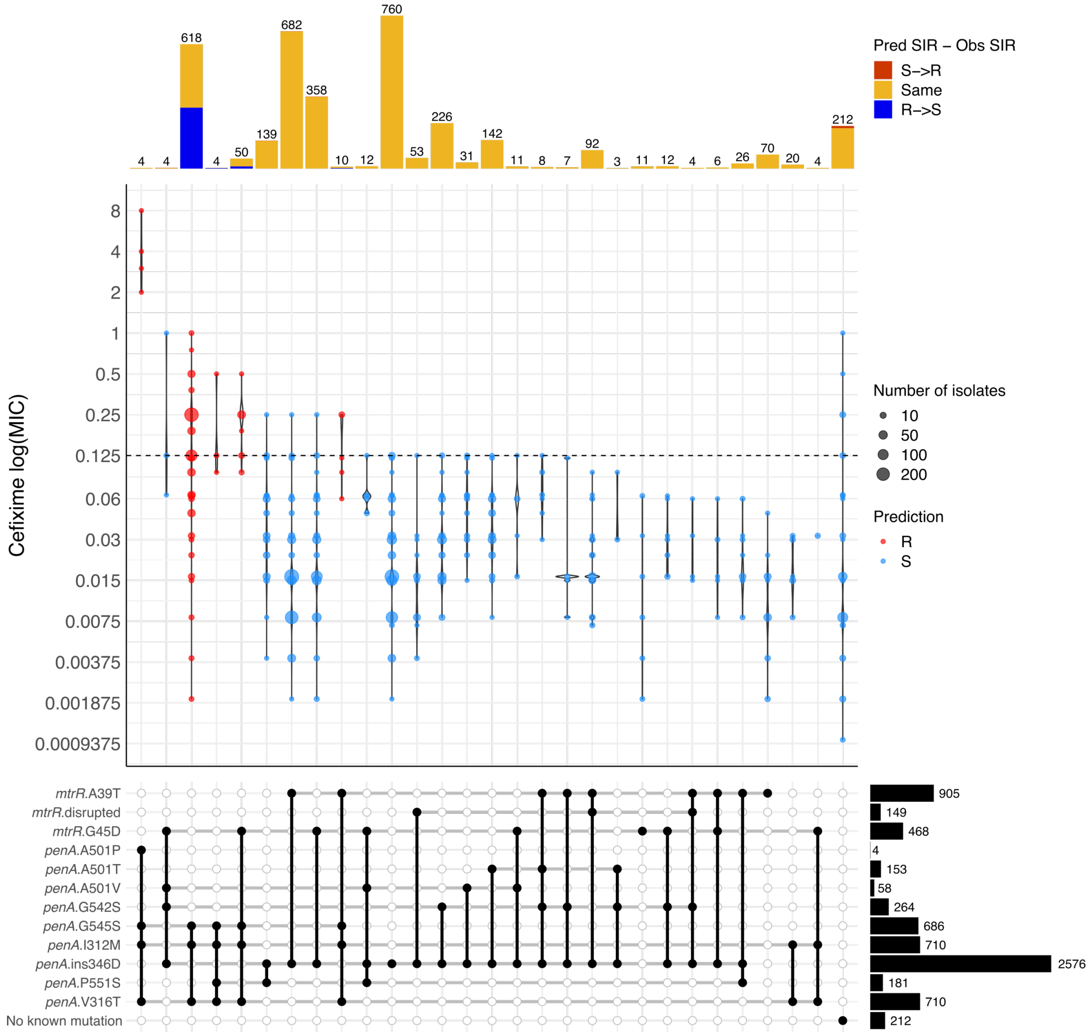


Figure S7. Distribution of minimum inhibitory concentration (MIC) (mg/L) values for cefixime in a collection of 3,987 *N. gonorrhoeae* isolates with different combinations of antimicrobial resistance (AMR) genetic mechanisms. Combinations observed in at least 3 isolates are shown. Dashed horizontal lines on the violin plots mark the EUCAST resistance breakpoint. Point colours inside violins represent the genotypic AMR prediction by Pathogenwatch on each combination of mechanisms. Barplots on the top show the abundance of isolates with each combination of mechanisms. Bar colours represent the differences between the predicted (Pred SIR) and the observed SIR (Obs SIR) (i.e. red for a predicted susceptible mechanism when the observed phenotype is resistant). Black bars on the bottom right represent the number of isolates with each particular mechanism individually or in combination.

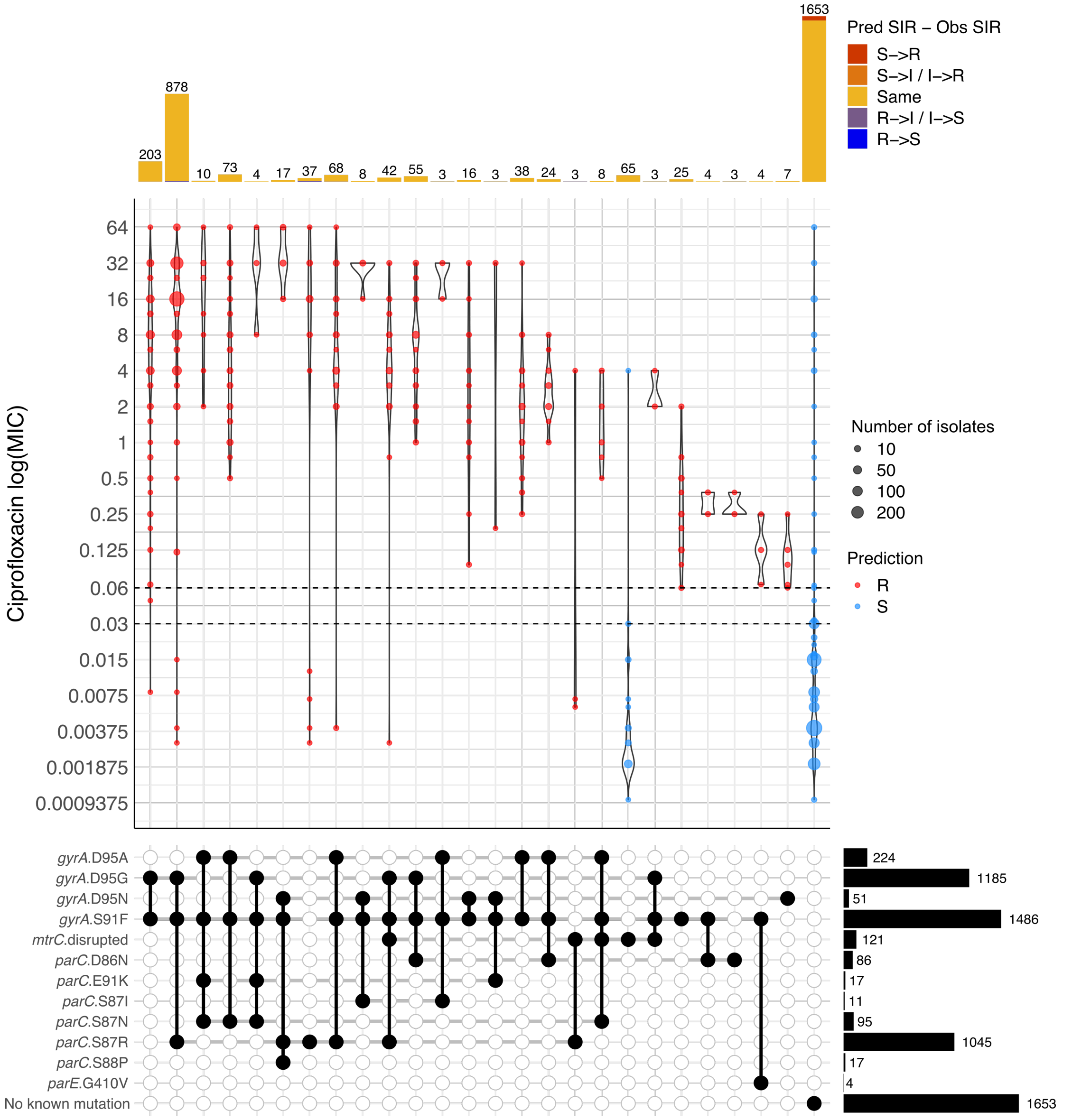


Figure S8. Distribution of minimum inhibitory concentration (MIC) (mg/L) values for ciprofloxacin in a collection of 3,987 *N. gonorrhoeae* isolates with different combinations of antimicrobial resistance (AMR) genetic mechanisms. Combinations observed in at least 3 isolates are shown. Dashed horizontal lines on the violin plots mark the EUCAST breakpoints. Point colours inside violins represent the genotypic AMR prediction by Pathogenwatch on each combination of mechanisms. Barplots on the top show the abundance of isolates with each combination of mechanisms. Bar colours represent the differences between the predicted (Pred SIR) and the observed SIR (Obs SIR) (i.e. red for a predicted susceptible mechanism when the observed phenotype is resistant). Black bars on the bottom right represent the number of isolates with each particular mechanism individually or in combination.

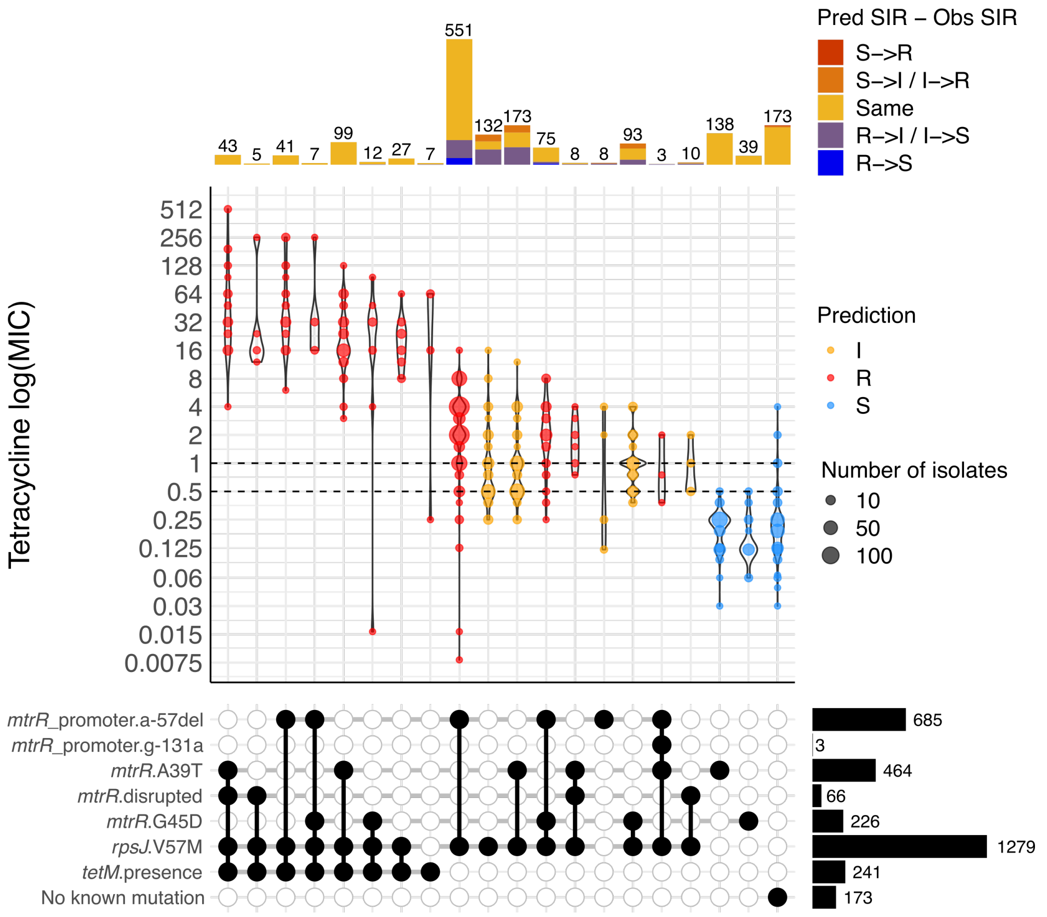


Figure S9. Distribution of minimum inhibitory concentration (MIC) (mg/L) values for tetracycline in a collection of 3,987 *N. gonorrhoeae* isolates with different combinations of antimicrobial resistance (AMR) genetic mechanisms. Combinations observed in at least 3 isolates are shown. Dashed horizontal lines on the violin plots mark the EUCAST breakpoints. Point colours inside violins represent the genotypic AMR prediction by Pathogenwatch on each combination of mechanisms. Barplots on the top show the abundance of isolates with each combination of mechanisms. Bar colours represent the differences between the predicted (Pred SIR) and the observed SIR (Obs SIR) (i.e. red for a predicted susceptible mechanism when the observed phenotype is resistant). Black bars on the bottom right represent the number of isolates with each particular mechanism individually or in combination.

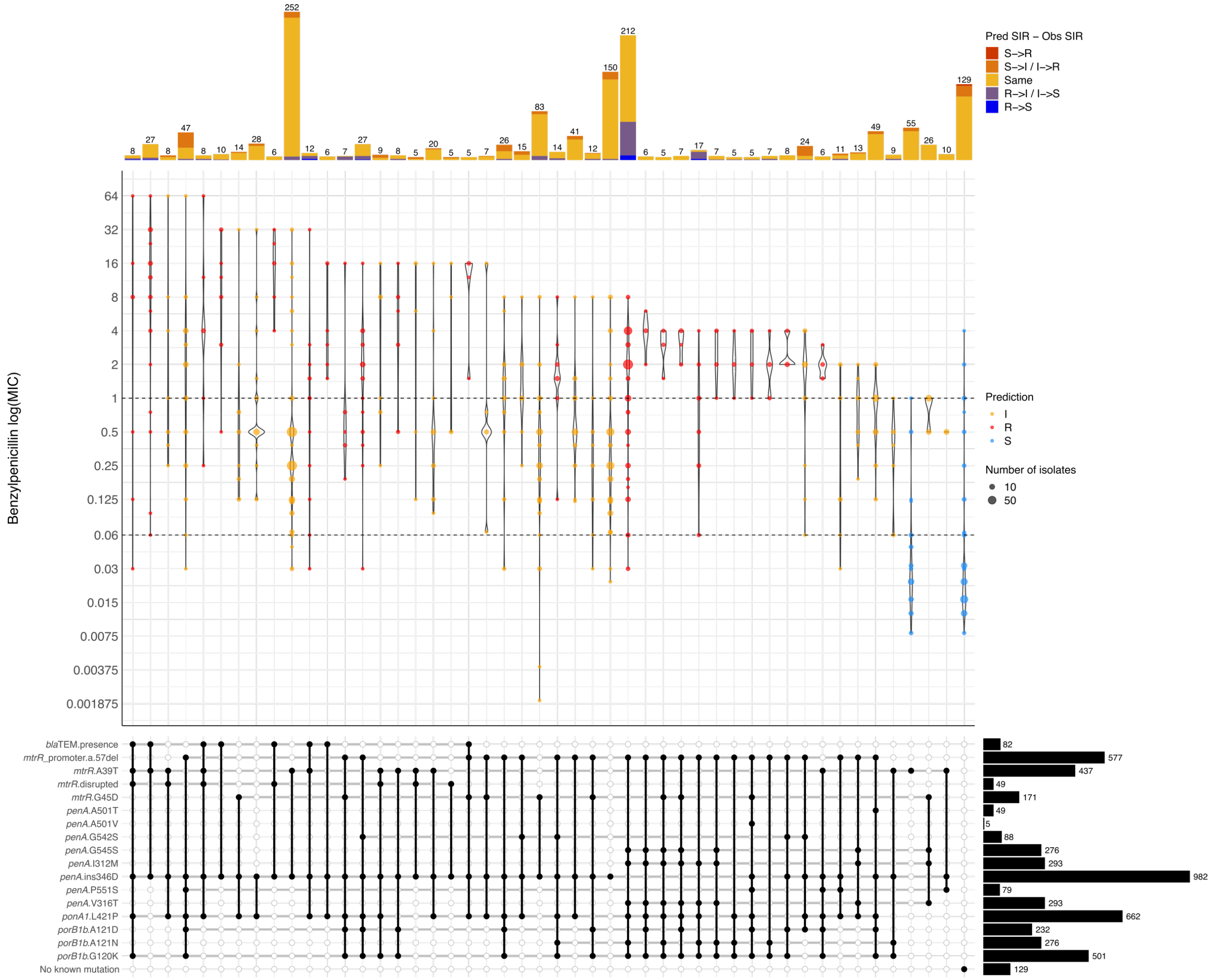


Figure S10. Distribution of minimum inhibitory concentration (MIC) (mg/L) values for benzylpenicillin in a collection of 3,987 *N. gonorrhoeae* isolates with different combinations of antimicrobial resistance (AMR) genetic mechanisms. Combinations observed in at least 5 isolates are shown. Dashed horizontal lines on the violin plots mark the EUCAST breakpoints. Point colours inside violins represent the genotypic AMR prediction by Pathogenwatch on each combination of mechanisms. Barplots on the top show the abundance of isolates with each combination of mechanisms. Bar colours represent the differences between the predicted and the observed SIR (i.e. red for a predicted susceptible mechanism when the observed phenotype is resistant). Black bars on the bottom right represent the number of isolates with each particular mechanism individually or in combination.

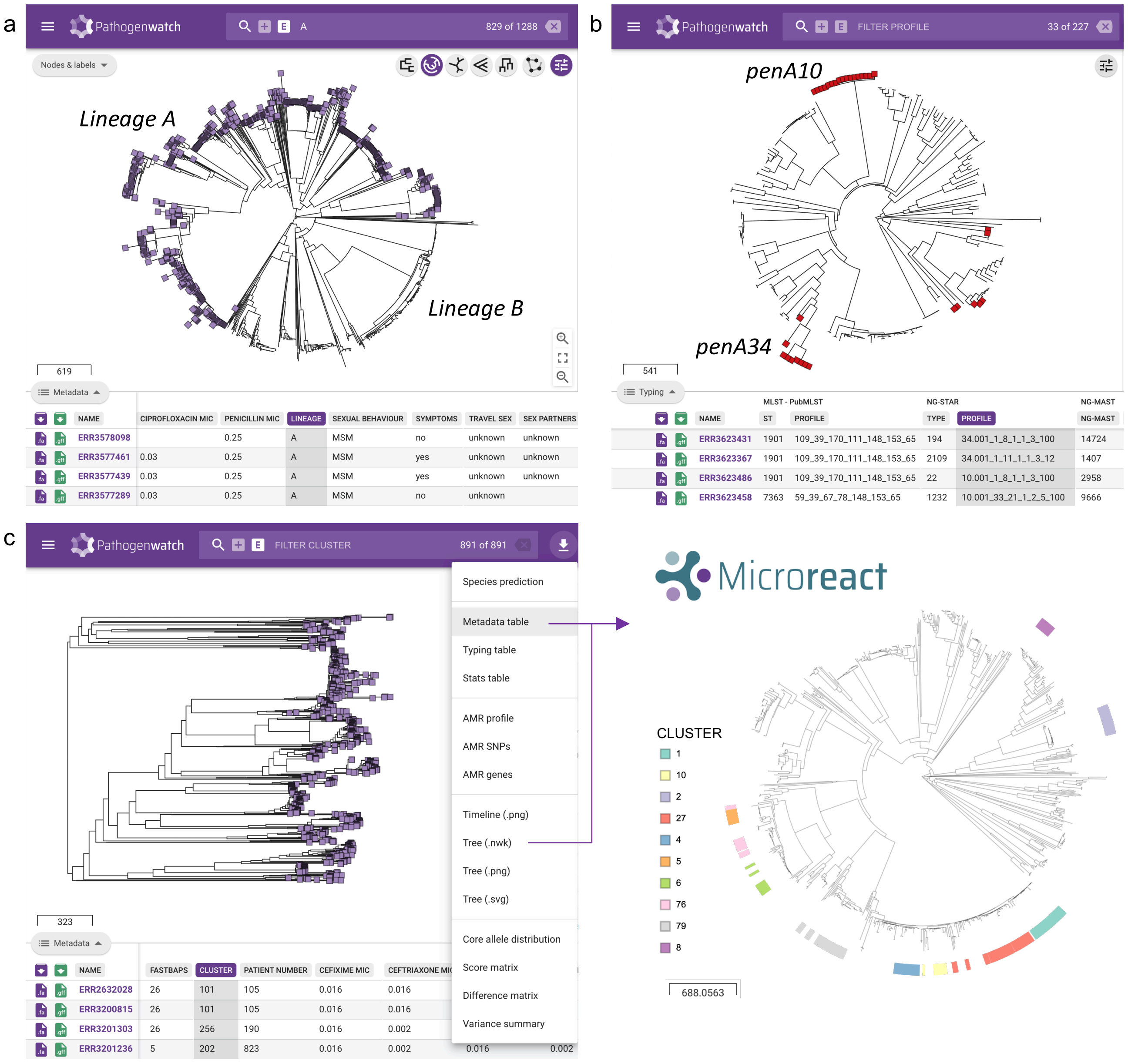

Figure S11. Major *N. gonorrhoeae* population lineages and clusters observed in scientific publications are consistent with the Pathogenwatch phylogenetic trees built from the same data obtained from public repositories. These examples show (a) two major lineages A and B from England reported in Town *et al* (2020) (68, 117), (b) two cefixime-resistant clades carrying *penA10* and *penA34* mosaic sequences in Vietnam reported in Lan *et al* (2020) (124) and (c) ten major clusters observed circulating in New York City as reported in Mortimer *et al* (2020) (120). In (c), the metadata table and phylogenetic tree was downloaded from Pathogenwatch and uploaded to Microreact for a more detailed visualization.

a

b

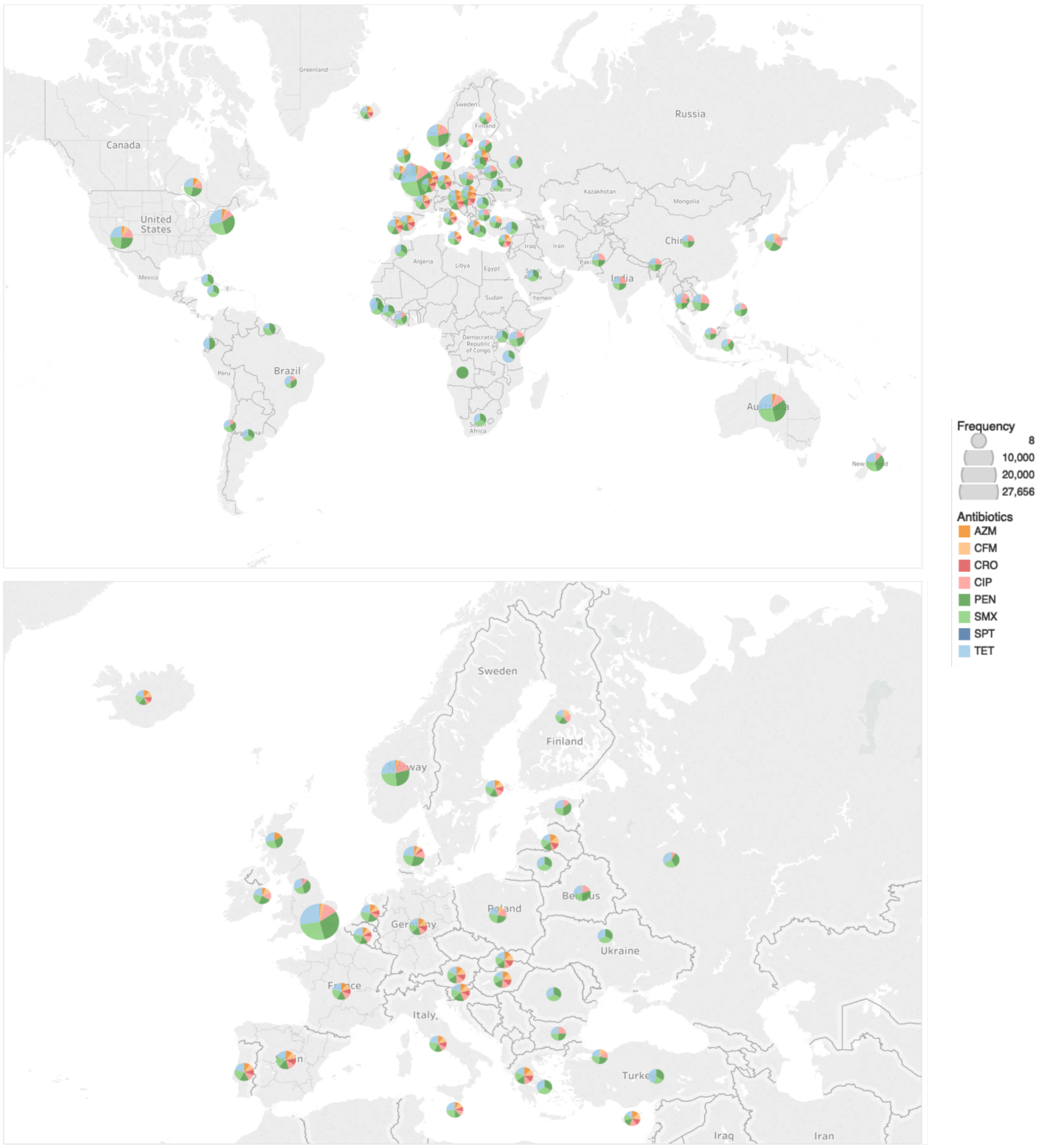


Figure S12. Geographical distribution of public *N. gonorrhoeae* genomes in Pathogenwatch from 27 different studies (Additional file 1: Table S6) worldwide (a) and Europe (b). The size of the pie charts is proportional to the number of isolates per country and the colours represent predicted antimicrobial resistance (AMR) to 8 different antibiotics using Pathogewatch AMR. AZM = Azithromycin, CIP = Ciprofloxacin, CFM = Cefixime, CRO = Ceftriaxone, PEN = Benzylpenicillin, TET = Tetracycline, SPT = Spectinomycin. Maps were represented using Tableau Desktop.
